# The influence of internal pressure and neuromuscular agents on *C. elegans* biomechanics: an empirical and multi-compartmental *in silico* modelling study

**DOI:** 10.1101/2023.11.09.566289

**Authors:** Clara L. Essmann, Muna Elmi, Christoforos Rekatsinas, Nikolaos Chrysochoidis, Michael Shaw, Vijay Pawar, Mandayam A. Srinivasan, Vasileios Vavourakis

**Affiliations:** Department of Bioinformatics and Molecular Genetics, University of Freiburg, Baden-Wuerttemberg, Germany; Department of Computer Science, University College London, Gower Street, London, UK; Software and Knowledge Engineering Lab, NCSR “Demokritos”, Athens, Greece; Department of Mechanical Engineering and Aeronautics, University of Patras, Rion-Patras, Greece; National Physical Laboratory, Hampton Road, Teddington, United Kingdom; Department of Mechanical and Manufacturing Engineering, University of Cyprus, Nicosia, Cyprus; Department of Medical Physics and Biomedical Engineering, University College London, London, United Kingdom

**Author notes:** Correspondence: C.L.E.: Department of Bioinformatics and Molecular Genetics, Institute of Biology III, University of Freiburg, Baden-Wuerttemberg, 79104, Germany; V.V.: Department of Medical Physics and Biomedical Engineering, University College London, London WC1E 6BT, United Kingdom.

**Keywords:** Caenorhabditis elegans, finite element method, biomechanics, atomic force microscopy, Aldicarb, osmotic shock, optogenetics

## Abstract

The function of a specific tissue and its biomechanics are interdepended, with pathologies or ageing often being intertwined with structural decline. The biomechanics of Caenorhabditis elegans (*C. elegans*), a model organism widely used in pharmacological and ageing research, has been established as biomarker for healthy ageing, though the mechanics of individual tissues have remained elusive. In this study we investigated the biomechanics of healthy *C. elegans* cuticle, muscle tissue, and pseudocoelom using a combination of indentation experiments and *in silico* modelling. Nematode stiffness measurements were performed using an atomic force microscope. The worm’s cylindrical body was approximated using a novel three-compartmental nonlinear finite element model, enabling analysis of how changes in the elasticity of individual compartments affect the bulk stiffness of *C. elegans*. The parameters of the model were then fine-tuned to match the simulation force-indentation output to the experimental data. To test the finite element model, distinct compartments were modified experimentally. Our *in silico* results, in agreement with previous studies, suggest that hyperosmotic shock reduced stiffness by decreasing the *C. elegans*’ internal pressure. Unexpectedly, treatment with the neuromuscular agent aldicarb, traditionally associated with muscle contraction, reduced stiffness by decreasing the internal pressure. It challenges previous assumptions about the effects of aldicarb. Furthermore, our finite element model can offer insights into how drugs, mutations or processes like ageing target individual tissues.

## 1 INTRODUCTION

The nematode Caenorhabditis elegans (*C. elegans*) is a powerful model organism for the study of different physiological processes including neuronal signalling, stress response and ageing, all of which ultimately converge into behavioural changes. Moreover, nematode behaviour has been extensively studied to test or investigate drugs, and to analyse the effects of genetic mutations. Here, automated, multidimensional *C. elegans* motion trackers paired with machine learning have generated databases for behavioural phenotypes and tracked life and health span effects of genetic and pharmacological interventions (Stroustrup et al., 2013; Yemini et al., 2013; Javer et al., 2018).

In the context of behavioural responses, locomotion plays an essential role. The worm’s locomotion involves a complex interplay between its neuronal network, contracting muscles, and the surface traction between the cuticle and the culture medium. Notably, locomotion also depends on the biomechanics of the worm’s tissues. As the worm ages, its body gradually becomes softer and loses the ability to move. We have previously analysed the biomechanics of ageing worms by atomic force microscopy and revealed a correlation between loss of stiffness and diminished mobility, establishing stiffness as a reliable biomarker for healthy ageing (Essmann et al., 2020). Since our stiffness calculations were based on the well-established Hertzian model for contact mechanics (Sneddon, 1965) by assuming the worm’s body being a single uniform tissue, it resulted in one bulk stiffness value for the entire worm indifferent to individual tissue mechanics (Essmann et al., 2020). Ultimately, and to differentiate between the decay of individual tissues, tissue properties are required to be mapped for each tissue layer of the *C. elegans* individually.

Like other organisms, *C. elegans* is composed of distinct tissue layers, including the cuticle, hypodermis, muscles and the pseudocoelom. The latter being a fluid filled cavity enclosing internal organs, which contributes to the overall biomechanics and structural support of the worm through hydrostatic pressure, as documented in [WORMATLAS](https://www.wormatlas.org/hermaphrodite/pericellular/mainframe.htm). Each *C. elegans* tissue layer is unique in its molecular and cellular composition thus likely different in stiffness, but very little is known about the local material properties and stiffness of these individual layers, and how they contribute to the overall body stiffness. Determining the stiffness values for each layer becomes crucial when examining the impact of specific mutations (e.g., mutations affecting muscle function or the extracellular matrix) external influences (food, temperature, salt content of the media) or processes such as ageing on specific tissues. This lack of knowledge partly arises from the challenge of directly measuring the deeper layers using current *in vivo* experimentation procedures. Recently, researchers analysed the stiffness of the dissected worm cuticle and how it is affected by collagen mutations or ageing using tensile test techniques (Rahimi et al., 2022). However, the cuticle alone does not determine the mechanics of the worm. Deeper layers, such as the muscle layer and hydrostatic pressure, have been experimentally modified to analyse their contribution to the overall stiffness of the worm, albeit their individual stiffness values have not been determined (Park et al., 2007; Petzold et al., 2011; Backholm et al., 2013). Petzold and his colleagues used piezoresistive cantilevers to find that chemic- or optogenetic-induced muscle contractions significantly increased the bulk stiffness of *C. elegans*, while muscle relaxation reduced it. Moreover, Park et al. described that puncturing the worm with a fine needle to release the hydrostatic pressure had modest effect on the overall biomechanics of the worm. One approach to overcome these limitations is to use computational modelling and to develop a new biomechanical model capable of assigning mechanical properties to distinct individual layers.

The biomechanics of *C. elegans* have been modelled in various ways based on experimental interrogations as early as 2005 in (Park et al., 2005, 2007) and later in (Nakajima et al., 2009). These studies incorporated a custom-designed MEM device and an environmental-scanning electron microscope (E-SEM) to measure *C. elegans* stiffness at the microscopic level. Utilising experimentally measured force-displacement (F-D – indentation depth) data, Park et al. employed a linearized Hertzian contact model, whereas Nakajima et al. employed a Euler-buckling theory model to estimate nematode bulk tissue stiffness. Further, Gilpin et al. (2015) employed a linear-spring mechanical analogue to explicitly model hydrostatic pressure, by taking the difference between the external and the desired internal pressure, to explore *C. elegans* whole-body biomechanics. A previous study in our lab by Elmi et al. (2017) introduced a three-dimensional finite element (FE) model of the *C. elegans*, assigning distinct stiffness values to an outer (cuticle, hypodermis, muscle) and an inner layer (pseudocoelom). This model was based on linear-elastic isotropic, homogeneous soft matter. More recently, Sanzeni et al. (2019) modelled *C. elegans*’ body as a tapering cylinder that consists of an outer shell structure (i.e., the cuticle, hypodermis, and body wall muscles) and an inner (tube) structure of the pseudocoelom with the intestine and gonad gland. To simplify their structural model, they employed a Hookean stress/strain constitutive law for the worm (thin shell) solid body. A more advanced model that uses a second-order elasticity theory to capture larger amplitude deformations and material nonlinearity was proposed by Du et al. (2023).

This study combines a new experimental procedure based on atomic force microscopy with the development of a new multi-compartmental FE model. We demonstrate the importance of internal pressure as the primary determinant of overall body stiffness and that hyperosmotic shock significantly reduces body stiffness. Moreover, our data show that the neuromuscular agents aldicarb and tetramisole reduce the internal pressure within the worm. This has important implications for our understanding of the effects of neuromuscular agents and their effect on the biomechanics of *C. elegans*. Moreover, the model can be used in the future to predict how mutations or pharmacological interventions impact the age-related decline of individual tissues.

## 2 MATERIALS AND METHODS

### 2.1 *C. elegans* laboratory maintenance

*C. elegans* wild-type (N2) and ZX299 animals were obtained from the Caenorhabditis Genetics Centre. ZX299 are genetic mutants lin-15B&lin-15A(n765) X with the extrachromosomal expression construct zxEx22[myo-3p::ChR2(H134R)::YFP+lin-15(+)] expressing channel rhodopsin in the body wall muscles. All animals were maintained using standard normal growth conditions and procedures (Brenner, 1974). For experimental purposes L4 animals were grown at 20°C until reaching the adult stage.

### 2.2 Drugs and osmotic shock treatments

Prior to the force-displacement measurements the worms were treated for 60 min with 15 mg/ml BDM (2,3-butanedione monoxime, Sigma-Aldrich), or 1 mM aldicarb (Sigma-Aldrich), or for 90–120 min with 100 *µ*M tetramisole (Sigma-Aldrich) in solution until they were no longer moving. For acute hyperosmotic shock treatment, worms were incubated in 500 mM NaCl in solution for one hour before force measurement containing 15 mg/ml BDM to prevent movement during measurements. For comparison between drugs in solution versus plate, worms were treated with 15 mg/ml BDM in solution or on NGM plate, or 1 mM aldicarb in solution or on NGM plate for 60min. For morphology analysis, worms were treated with either the drug solution or hyperosmotic solution (500 mM NaCl) at the same concentrations as described above. For the optogenetics: ZX299 transgenic worms were cultivated on agar plates with OP50 bacteria in the presence of all-trans retinal (Sigma-Aldrich) (Nagel et al., 2005). Worms were treated as described above prior to the force-measurements. For stimulation of channelrhodopsin, worms were illuminated with 450–490 nm light for defined time to measure force in response to change in body stiffness.

### 2.3 Force measurements

#### 2.3.1 Atomic Force Microscopy (AFM)

Treated worms were transferred and glued to a 1 mm thick 4% agarose bed in a petri dish as described previously (Essmann et al., 2017). Force measurements were taken from the neck and hip region of the worm, avoiding the mid body or vulva region, see (Essmann et al., 2020). Preparation of the worms and AFM measurements were all performed at RT. Individual force-displacement curves were acquired using a NanoWizard3 AFM (JPK) in force spectroscopy mode (set force 450 nN; 0.5 mm/s indentation speed). All data were captured using a 10 *µ*m diameter glass bead attached to a tipless cantilever of k = 5.79–10.81 N/m stiffness (NSC12 7.5 N/m *µ* Masch produced by [sQUBE](http://www.sqube.de/)) to prevent the cuticle from being pierced at larger indentations. Cantilever sensitivity and spring constant were calibrated using the JPK calibration tool (thermal noise method (Butt and Jaschke, 1995)) prior to each experiment.

#### 2.3.2 Micro-Force Displacement System (µFDS)

For the larger amplitude force-displacement experiments we used an in-house customized micromanipulation setup to allow uniaxial indentation of *C. elegans* using a microforce sensing probe (Elmi et al., 2017). Using cover glass with a 2% agarose pad, the drugs treated immobile worms were glued on to the edge of a second coverslip using Dermabond glue (2-octyl cyanoacrylate, Suturonline.com) and subsequently immersed in M9 buffer. The worm sample was placed in an upright widefield fluorescence microscope (BX51WI, Olympus) with 20x/1.0 water immersion objective lens (LUMPlanFL N, Olympus) fluorescence filter cube (Semrock) and a scientific CMOS camera (Orca-Flash4.0 v2, Hamamatsu Photonics). The body of each worm was indented using microforce sensing probe (FT-S100, FemtoTools) fitted with tungsten tip. The position of the probe was controlled using a motorized 4-axis stage system (ECS series, Attocube), which allowed precise positioning of the tip within (x, z) and perpendicular (y) to the focal plane of the microscope, as well as adjustment of the in-plane tilt. Animals were mounted on a separate kinematic stage system decoupled from the microscope body and the probe.

#### 2.3.3 AFM and µFDS data analysis

Raw AFM data were analysed using JPK analysis software. All individual force curves were processed to zero the baseline, to determine tip-sample contact point and to subtract displacement of the tip due to cantilever bending. To calculate the Young’s Modulus, the mean, minimum and maximum force-indentation curve (Fig 1d) was further analysed by fitting the Hertz/Sneddon model for contact mechanics to the entire curve by using the JPK software and by taking the indenter shape (10 *µ*m diameter bead) into account (see Fig 1f). The µFDS directly recorded the force data as displayed in the graphs with minimal processing: The recording of the force probe movement towards the sample was cut off, and force and displacement zeroed just before touching the worm (confirmed optically).

**Figure 1.**
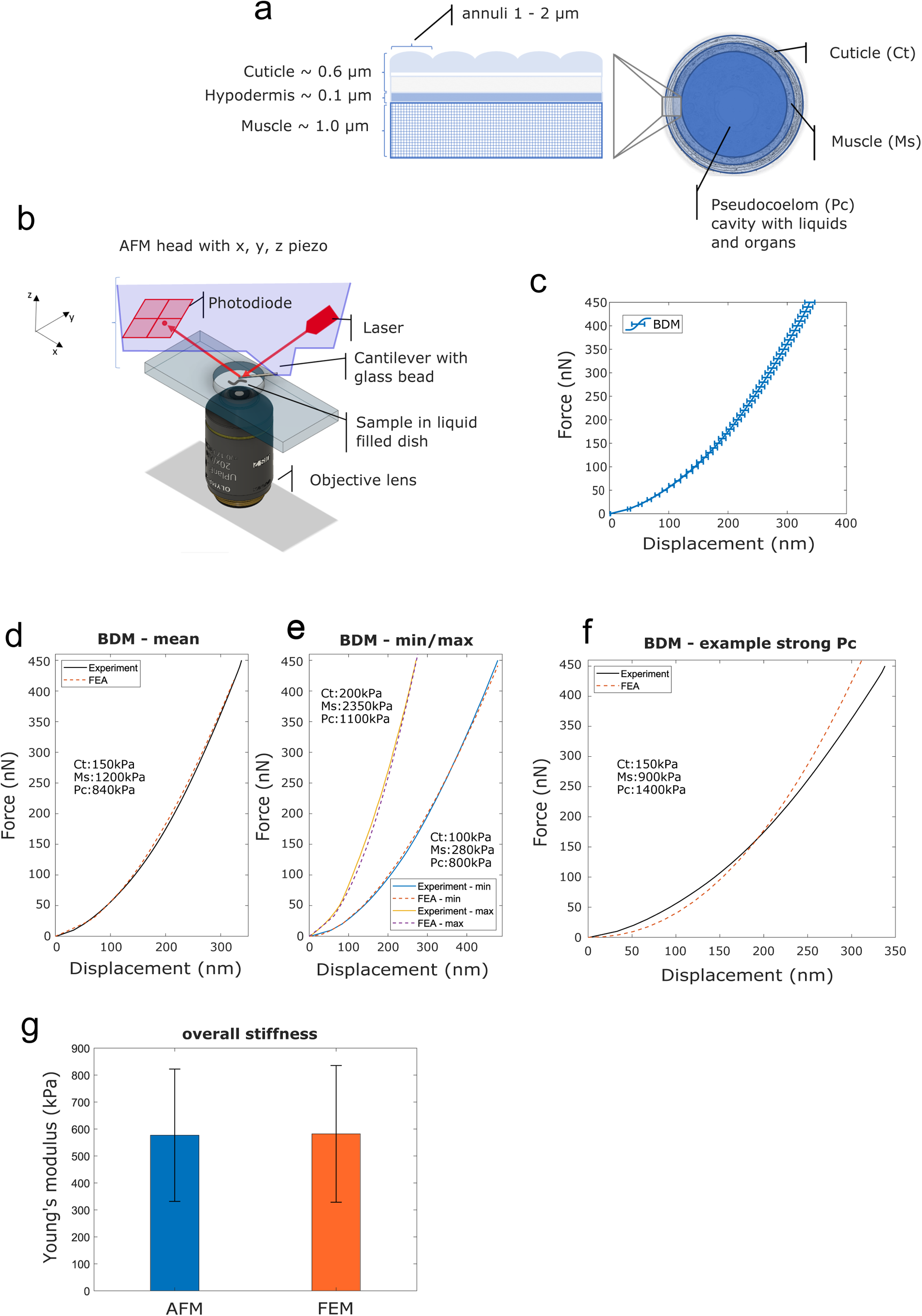
**(a)** Illustration of *C. elegans*’ body composition – structure of the outer layers (cuticle, hypodermis and muscle tissue) (left) and a cross sectional view (right). **(b)** Schematic drawing of the AFM probing device setup. **(c)** Mean experimental F-D curve for *C. elegans*’ treated with BDM, measured using AFM. Horizontal bars denote standard error from the mean. **(d, e)** Experimental (AFM) and simulated (FE model) F-D response curves corresponding to nematodes with mean, the minimum / maximum stiffness respectively. **(f)** Experimental data (BDM control) and simulated F-D response curve from the FE model with decreased Young’s modulus for the muscle component and increased Young’s modulus for the pseudocoelom. **(g)** Overall Young’s modulus of *C. elegans* – comparison between AFM-software model (Hertz-Sneddon) and FE model simulation. Vertical error bars denote standard deviation from the mean.

To calculate the nematode bulk stiffness from either the AFM and or the µFDS force-displacement data, linear regression was applied on the data within the [0.5 *δ_max_*, *δ_max_*] range, where *δ_max_* the maximum displacement value.

#### 2.3.4 Imaging and analysis

Body measurements: Images were acquired using a CMOS camera (Orca-Flash4.0 v2 Hamamatsu Photonics). The ImageJ plugin WormSizer was used for detecting the worm size, measuring body length and diameter, and body volume calculated according to Jung et al. (2012) and Moore et al. (2013).

### 2.4 *C. elegans* FE model

To realistically simulate nematode biomechanics, an advanced *C. elegans* finite element (FE) model was developed using the proprietary software ABAQUS/Standard (SIMULIA, 2014). The proposed 3D FE model overcomes two major simplifications of published biomechanical models: (*a*) it distinguishes the major tissue regions of *C. elegans* that primarily contribute to the worm’s body biomechanics, which are modelled individually (with separate model parameters). In addition, (*b*) our FE model accommodates nonlinear (material model) biomechanical behaviour of the tissue of *C. elegans* and the nonlinearities that appear from the mechanical interactions with components of the testing device (i.e., the AFM tip). Part of the nematode body was approximated in ABAQUS as an idealized cylinder (see Suppl Text 1) subdivided into three distinct tissue compartments corresponding to the cuticle and dermis, the muscle tissue, and the pseudocoelom (gonad, intestine, etc.). Each tissue compartment was modelled using separate material properties (i.e., stiffness) based on a Green-elastic, neo-Hookean constitutive model (see Suppl Text 1). Next, for the FE discretization of the *C. elegans* model, a hexa-dominant 3D mesh was generated in ABAQUS, with a finer mesh used close to the region where the tip contacts the cuticle, growing further away from the indentation region. A mesh sensitivity and convergence analysis found the optimal FE mesh had a 0.4 *µ*m minimum edge size. Finally, proper contact and boundary conditions were defined to reduce the size of the computational domain and replicate the constraints applied to the nematode movement *in vivo* during testing (see Suppl Text 1). Computer simulations were run on a Dell workstation with an Intel(R) Core(TM) i7-6800K CPU (3.40GHz) and 64 GB of RAM, with each *in silico* experiment taking up to fifteen minutes.

## 3 RESULTS

### 3.1 Three-compartment nonlinear biomechanical FE model of *C. elegans*

The *C. elegans* body is modelled using the FE method, idealized as a cylindrical structure, which consists of four distinct compartments; the cuticle (0.6 *µ*m) is the outermost layer that serves as an exoskeleton, the hypodermis (0.1 *µ*m) a very thin cellular layer, the muscle tissue (1.0 *µ*m), and the pseudocoelom, a fluid filled cavity with internal organs and gonads (Lints and Hall, 2005; Wolkow and Hall, 2016; Altun and Hall, 2023), see Fig 1a. Previously we had used a simple two layered model and large force-displacement (F-D) measures of up to 14 *µ*m to describe *C. elegans* biomechanics (Elmi et al., 2017); thus, combining the three outer layers into one compartment. In this study, we wanted to elucidate on the role of the distinct components of this previously named ‘outer’ layer and each of their contribution to the overall stiffness. We therefore required a force sensing system in the range of nanonewtons that is capable of indentations in the range of nanometres to a few micrometres. For this purpose, we used an atomic force microscope (AFM; Fig 1b), to produce data to inform and validate the FE model used to simulate the *C. elegans* biomechanics (see also Suppl Text 1 and in paragraph ‘*C. elegans* FE model’ above).

As in our previous studies (Elmi et al., 2017; Essmann et al., 2017), we used 2,3-butanedione monoxime (BDM, a chemical that has been described to relax muscles (Petzold et al., 2011; Backholm et al., 2013) by inhibiting myosin (Higuchi and Takemori, 1989; Soeno et al., 1999)) to immobilise worms during AFM analysis. Using a 10 *µ*m bead attached to a tipless cantilever to avoid damaging or penetrating the cuticle during the indentation process, we applied a set force of 450 nN and recorded individual force-indentation curves. The mean indentation at this force was 338 nm *±* 8.4 SEM (n=44; Fig 1c). Subsequently we employed our FE model to simulate the AFM indentation test and to reproduce the F-D plots obtained from the experiments. Following an Edisonian approach (Wills, 2019; Dandekar et al., 2003), we attempted through the simulations to interrogate and estimate the model parameter values (Young’s modulus) for each of the three tissue compartments of the FE model: the cuticle (cuticle and hypodermis), the muscle, and the pseudocoelom. Fig 1d shows the mean force (nN) versus the displacement (nm) curve of the experiments (black solid line) compared against the FE-simulated curve (red-dashed line). After optimising for the Young’s modulus of each of the three compartments of the *C. elegans* FE model, we converged to the following set of values that gave the best agreement to the experiments: 150 kPa, 1200 kPa and 840 kPa for the cuticle, muscle and pseudocoelom respectively (Fig 1d and Suppl Fig 1a). To explore the range of natural variation between worms’ biomechanics, we attempted to reproduce the two extreme F-D curves of the experiments (min, max) with the FE model, and quantified the difference in Young’s modulus for each tissue compartment (Fig 1e). Taking the natural variation into consideration, the mechanical properties we propose for BDM-treated wild type animals range from 200 kPa to 100 kPa for the cuticle, from 2350 kPa to 280 kPa for the muscle layer and from 1100 kPa to 800 kPa for the pseudocoelom (mean values: 150 kPa for the cuticle, 1200 kPa for the muscle, 840 kPa for the pseudocoelom).

Fig 1f shows a simulated F-D generated from the FE model with Young’s modulus values for the muscle tissue set to 900 kPa and for the pseudocoelom set to 1400 kPa. The nematode’s bulk stiffness predicted by the FE model is lower at the indentation range *<*150 nm and increases rapidly at indentation *>*200 nm, with poor agreement with the experimental F-D curve (solid line in Fig 1f). To validate our FE model of the *C. elegans*, we compared the numerically calculated overall Young’s modulus of *C. elegans* to that obtained from the Hertz/Sneddon model using the AFM-data analysis software. We found no significant difference (582 kPa *±* 253 kPa versus to 577 kPa *±* 245 kPa respectively) between the AFM measurements and FE model predictions, as shown in the bar plot of Fig 1g. For this, the overall stiffness values of the mean, min and max F-D curves were averaged (see Fig 1f).

### 3.2 Hydrostatic pressure contributes significantly to *C. elegans*’ bulk stiffness

To elucidate the contribution of the internal hydrostatic pressure to the stiffness of the worm, we exposed worms to a high salt buffer (0.5 M NaCl) to induce hyperosmotic osmotic shock and captured F-D curves using the AFM as previously. In high salt buffer exposed the set-point force of 450 nN was reached for a mean indentation depth of 1063 nm *±* 42 SEM (n=46). Under control conditions the same compressive force was reached at an indentation depth of 413 nm *±* 11 SEM (n=48), meaning that worms were significantly less stiff under hyperosmotic shock (Fig 2a). To quantify the difference in bulk stiffness between these two conditions, we employed linear regression on the data for moderate to high indentation depths, corresponding to probe displacement *>*200 nm for the BDM tests and *>*500 nm for high salt tests, and estimated the bulk stiffness of the nematode. The normalized values are seen in Fig 2b (for absolute values see Suppl Table 1).

**Figure 2.**
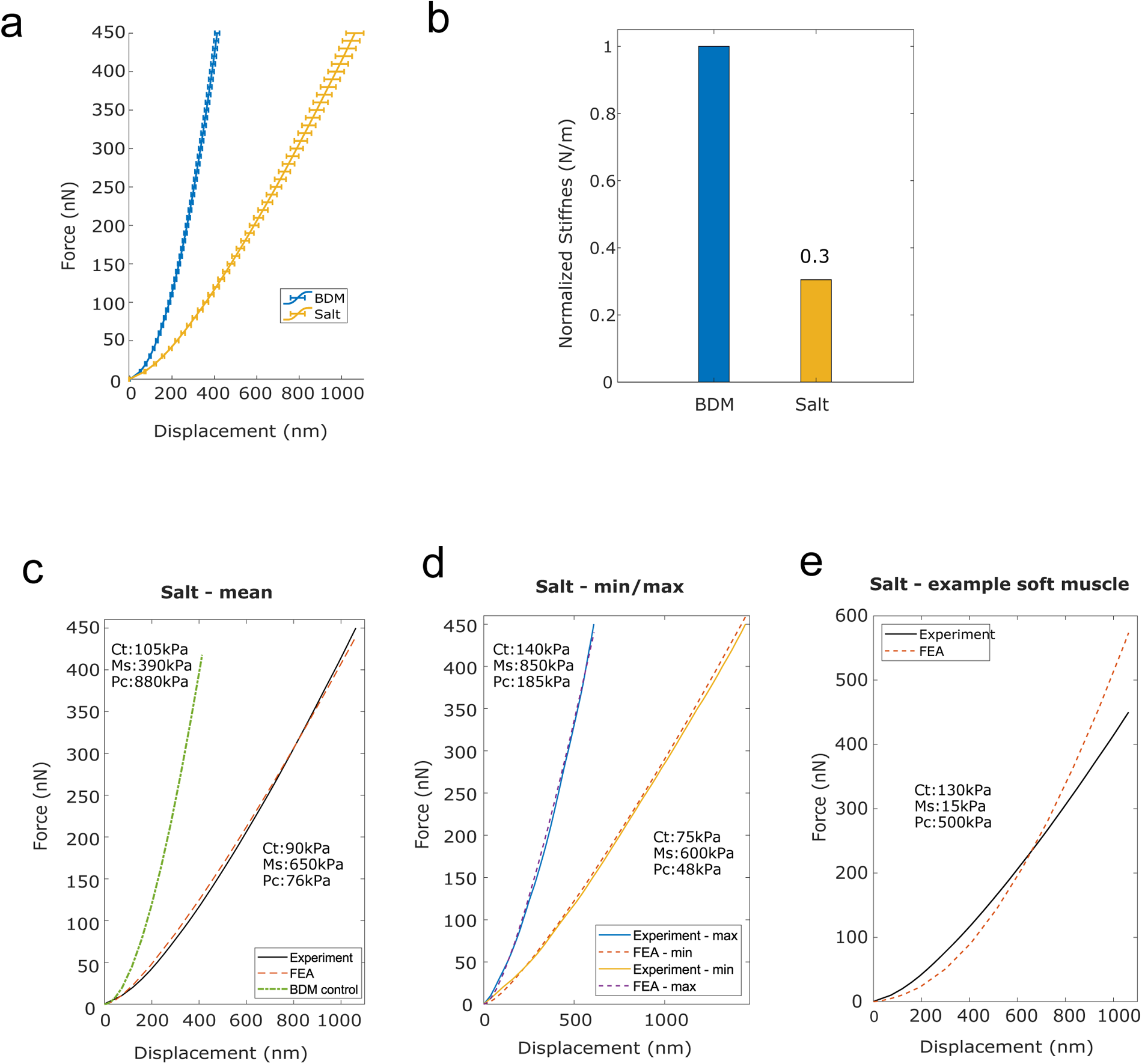
(**a**) Mean F-D curve for *C. elegans* treated with BDM alone (blue) and treated with BDM and exposed to high salt buffer (yellow), measured using AFM. Horizontal bars denote standard error from the mean. (**b**) Normalized bulk stiffness estimated from the F-D data. (**c, d**) Mean, and the minimum / maximum experimental (AFM) and simulated (FE model) F-D response curves for worms treated with high salt. (**e**) Experimental F-D curve for high salt treated worms and the F-D curve predicted by the FE model with a reduced value for Young’s modulus of the muscle component compared to the BDM treated control.

The Young’s modulus for each of the three compartments of the *C. elegans* FE model was then modified to fit the mean experimental F-D curve for high salt-treated worms (Fig 2c). After trial-and-error optimisation, the model parameters were: 90 kPa for the cuticle compartment, 650 kPa for the muscle compartment, and 76 kPa for the pseudocoelom. Comparing these values to the best fit parameters in to the control case (105 kPa, 390 kPa and 880 kPa; see green dash-dot line in Fig 2c and Suppl Fig 1b) we observe an 14% and a 91% approximately drop in the Young’s modulus of the cuticle tissue and the pseudocoelom respectively, whereas the muscle sees an *≈*67% increase of the Young’s modulus. In addition to the mean F-D curve on Fig 2c, we used the FE model to reproduce the two extreme F-D curves of the experiment (min, max), and again optimised the model to estimate Young’s Modulus for each tissue compartment. The simulations for the high salt-treated wild type *C. elegans* gave Young’s moduli that range within 140 kPa to 75 kPa for the cuticle, 850 kPa to 600 kPa for the muscle, and 185 kPa to 48 kPa for the pseudocoelom.

We then lowered the Young’s modulus for the muscle tissue and increased Young’s modulus for the pseudocoelom and re-ran the FE model. As shown in Fig 2e, the agreement of the numerically predicted F-D curve to the experimental data recorded under high salt experimental data was poor. This suggests that the hyperosmotic shock caused by salt had the largest impact on the pseudocoelom by reducing its pressure, attributed to loss of water due to osmosis, resulting in reduced volumetric resistance in that compartment. In addition, high salt treatment also increased muscle stiffness in line with previously reported results (Park et al., 2007). In conclusion, the overall worm body stiffness is significantly reduced in response to salt exposure.

### 3.3 Aldicarb treatment reduces *C. elegans*’ bulk stiffness

To understand the contribution of the body wall muscle layer to worm body stiffness we next modulated the muscle tone of the worm using the well-known chemical called aldicarb. Aldicarb has been described previously to hyper-contract muscles by blocking the degradation of acetylcholine at the neuro-muscular junction of *C. elegans* (Nurrish et al., 1999; Lackner et al., 1999). We treated the worms with either aldicarb or BDM as control condition until immobilized and measured their stiffness using the AFM (Fig 3a).

**Figure 3.**
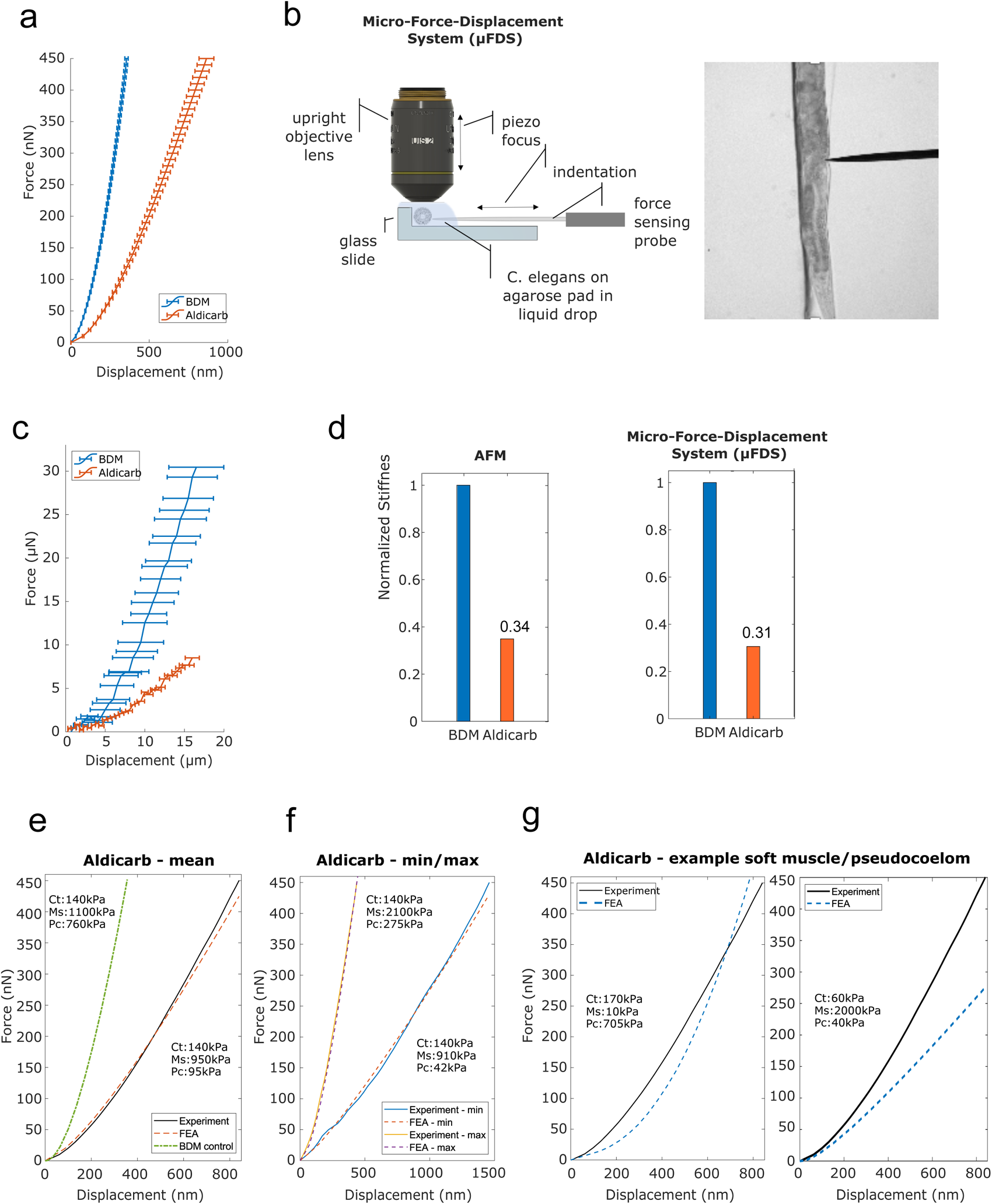
(**a**) Mean F-D data for *C. elegans* treated with BDM (*blue*) or aldicarb (*orange*) measured using AFM. Horizontal bars denote error deviation from the mean. (**b**) Schematic drawing of the micro-Force-Displacement System (µFDS) and representative brightfield micrograph showing a worm under indentation (right). (**c**) Mean F-D curves for *C. elegans* treated with BDM (*blue*) and aldicarb (*orange*) measured using the µFDS. Vertical bars denote standard error from the mean. (**d**) Normalized bulk stiffness estimated from the F-D curves measured using AFM (left) and µFDS (right). (**e, f**) Mean and minimum / maximum experimental (AFM) and simulated (FE model) F-D curves for worms treated with aldicarb. (**g**) Experimental F-D curve for aldicarb treated worms and the F-D curve predicted by the FE model with a reduced value for Young’s modulus of the muscle component (left) and the pseudocoelom compared to the BDM treated control.

The mean F-D curve of aldicarb-treated worms showed a maximum indentation of 864 *±* 47 nm (n=76) for a total compressive force of 450 nN, compared to 356 nm *±* 11 nm (n=68) for the BDM-treated worms. Contrary to our expectations, this result suggests that aldicarb-treated worms, although hyper-contracted, are softer than BDM-treated worms (Fig 3a). We hypothesized that when a worm shrinks due to muscle hypercontraction, the cuticle of the animal folds, thereby, allowing for a margin of the corresponding tissue to start becoming indented without significant compressive force being applied and therefore making the worm appear softer than in its uncontracted state. Based on this explanation, the stiffness should increase for larger indentations. To test this, we employed a Micro-F-D System (µFDS) as a second force-indentation tool (Fig 3b) able to indent worms up to 20 *µ*m (Elmi et al., 2017). BDM-treated worms were mounted on top of an agarose pad against the edge of a glass coverslip to provide a rigid surface when indenting the worm from the side with a uniaxial force sensor (Fig 3b). Using µFDS, we measured F-D curves for indentations of up to 16 *µ*m. For that indentation depth, we measured a mean compressive force of 29.3 *µ*N *±* 3.5 SEM (n=5) for *C. elegans* treated with BDM, whereas for when treated with aldicarb the same indentation resulted in a compressive force of only 8.5 *µ*N *±* 0.9 SEM (n=12; Fig 3c). Hence even at larger indentations, aldicarb treated worms are softer than BDM-treated worms. To quantify the difference in bulk stiffness between these two treatment conditions, and compare data from AFM and µFDS setups, we employed linear regression at moderate to high indentation depths to estimate the bulk stiffness using F-D data for indentations *>*10 *µ*m for the µFDS data set, and *>*200 nm for the BDM and *>*300 nm for the aldicarb AFM data set. The normalized values (Fig 3d; for absolute values see Suppl Table 1) indicate that the bulk stiffness of aldicarb-treated worms is decreased by 66% when compared to BDM-treated animals. To further investigate the effect of the neuromuscular agent, untreated worms were placed in the µFDS, and F-D data was captured over time after adding aldicarb. The results indicate a gradual decline in stiffness to reaching 40% of the initial value 150 minutes after treatment (Suppl Fig 5).

This unexpected result prompted us to investigate the effect of tetramisole on worm stiffness, another chemical known to activate cholinergic receptors on the body wall muscles and thereby cause muscles to hyper-contract over time (Lewis et al., 1980). Similarly, to the results for aldicarb-treated worms, data captured using AFM and µFDS systems for tetramisole-treated animals show they are significantly softer than BDM treated worms (see Suppl Fig 2).

We modified the Young’s modulus values for all three compartments of the FE model to reproduce the F-D curve for aldicarb-treated worms (Fig 3e). The numerically estimated Young’s modulus for the cuticle (140 kPa) and the muscle (950 kPa) compartments varied by less than 10% from the corresponding values for BDM-treated worms (Suppl Fig 1c). However, the estimated Young’s modulus for the pseudocoelom was 94 kPa, and dramatically lower than the 760 kPa for the pseudocoelom compartment of the BDM-treated worms. To also explore the range of natural variation between the aldicarb-treated worms’ biomechanics, we re-optimised the model to fit the two extreme experimental (min, max) F-D curves. This yielded estimates of between 42 kPa to 275 kPa for the Young’s modulus of the pseudocoelom, between 910 to 2100 kPa for the muscle tissue, whilst the Young’s modulus of the cuticle remained close to 140 kPa (Fig 3f). We also attempted to fit the experimental data by softening the muscle tissue and stiffening the pseudocoelom, or inversely by stiffening the muscle and softening the pseudocoelom as such. As seen in Fig 3g the simulated F-D curves (blue dotted lines) in both cases compare poorly to the mean experimental F-D curve for the aldicarb-treated worms (black solid line).

To conclude, rather than increasing stiffness treating *C. elegans* with aldicarb or tetramisole softens the worm. Fitting the experimental data using our multi-compartment FE model indicates that both agents significantly reduce the stiffness of the pseudocoelom without significantly changing the muscle layer compared to the control condition (i.e., BDM). This suggests that both these drugs have additional effects beyond their function as muscle contracting agents.

### 3.4 Osmotic shock and aldicarb treatment reduce worm size

Our FEM simulations indicated that worms treated with aldicarb loose most of their pseudocoelom stiffness. The reduced stiffness closely resembled that of worms exposed to an osmotic shock (Fig 2a) which causes loss of water and subsequently shrink in body size (Park et al., 2007). To understand more about the additional effects of aldicarb, we measured changes to the length, width and volume of the worms exposed to aldicarb or osmotic shock compared to BDM-treated controls (Fig 4a). Both osmotic shock and aldicarb treatment significantly decreased worm size with measured volumes of 2457.34 mm^3^ *±* 35.9 SEM and 2439.35 mm^3^ *±* 64.6 SEM respectively, compared to 2870.41 mm^3^ *±* 45.1 SEM for animals treated with BDM alone (n = 24, 49 and 67 respectively; Fig 4b). However, in contrast to worms subject to osmotic shock, aldicarb-treated worms reduced more in width (53.87 *±* 0.39 *µ*m, compared to 55.59 *±* 0.57 *µ*m) than length (1014.76 *±* 13.57 *µ*m compared to 907.22 *±* 9.22 *µ*m; Fig 4c and Fig 4d). It has been reported previously that worms exposed to aldicarb on plates shrink over time (Opperman and Chang, 1991; Miller et al., 1996; Nurrish et al., 1999; Lackner et al., 1999; Glenn et al., 2004; Mahoney et al., 2006). When we compared worms paralysed on aldicarb-containing plates versus aldicarb solution we found that animals exposed to aldicarb on plates shrank more with a reduced length and volume (Suppl Fig 3a). Based on analysis of AFM F-D curves these worms also tended to be 13% softer than worms paralysed in aldicarb solution (Suppl Fig 3b–3c). However, we observed no difference in stiffness or volume between worms treated with BDM in solution versus plate, although we found an *≈*10% difference in length (Suppl Fig 3d–3f).

**Figure 4.**
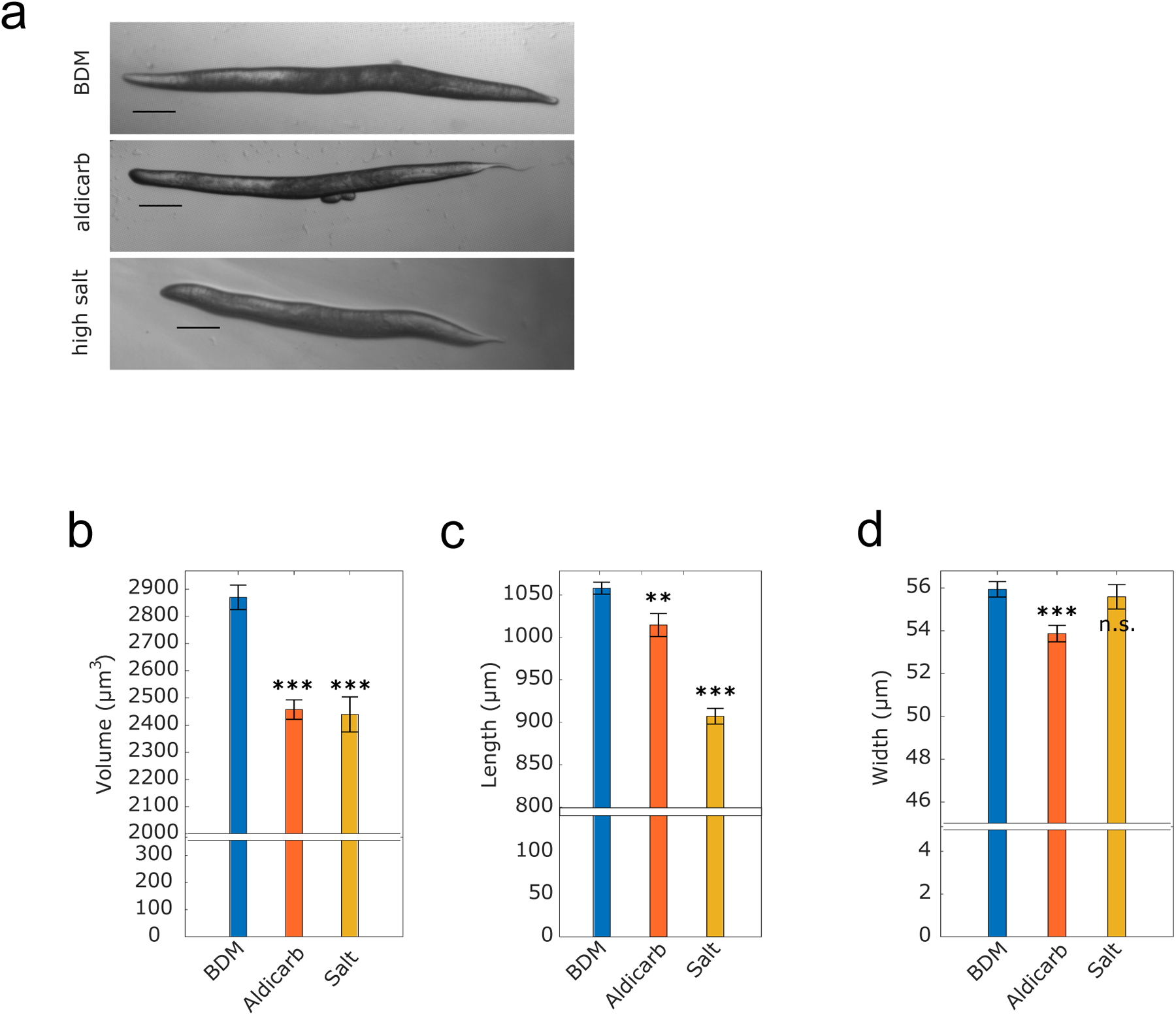
(**a**) Representative brightfield micrographs, ordered from top to bottom respectively, of worms paralysed in BDM, treated with aldicarb and high salt (scale bar: 100 *µ*m). (**b**) Mean volume, (**c**) mean length, (**d**) mean width of worms paralyzed with BDM (*blue*), aldicarb (*orange*) or high-salt (*yellow*) solution (n = 64, 49, and 24 respectively). Vertical bars denote standard error from the mean. P values are indicated as follows: n.s. = not significant, *∗∗ ≤* 0.01–0.001, and *∗ ∗ ∗ ≤* 0.001. Results were determined by two-tailed t-test.

### 3.5 Optogenetically controlled muscle contraction increases *C. elegans* bulk stiffness

Our data show that, consistent with previous reports (Mulcahy et al., 2013; Izquierdo et al., 2021), when aldicarb is administered in solution or on a plate the animals shrink. However, contrary to previous findings we measured a decrease in worm stiffness following aldicarb exposure (Park et al., 2007; Petzold et al., 2011). Moreover, our FEM simulations suggest this arises due to a change in the pseudocoelom rather than in the muscle layer (Fig 3e). It is possible that muscle contraction is not measurable with our force-indentation set-ups, or that chemically induced muscle contraction is accompanied by other effects and in combination this does not result in tissue stiffening. To investigate this possibility, we employed optogenetics as an alternative, non-chemical, tool to induce muscle contraction in *C. elegans*, using a worm strain expressing a channelrhodopsin-2 (ChR2) variant in its muscle tissue (Schultheis et al., 2011). ChR2 is an ion channel that responds to UV light by opening and allowing ions to enter the cell. When expressed in muscle cells ChR2 enables control of muscle contraction (AzimiHashemi et al., 2014). To test whether we could detect force associated with muscle contraction, we analysed untreated worms expressing channel rhodopsin using the µFDS setup. The AFM system was unsuitable for such measurements due to the uncontrolled, large movements of non-treated worms. The force sensor was used to indent the worm and clamp it against the coverslip edge. After a few seconds of baseline recordings, the worm was irradiated for 7–8 seconds with UV light to activate muscle contraction (Fig 5a). UV irradiation (On) resulted in a sharp rise in the compressive force sensed by the force sensing tip. When UV-light was turned off (Off) the force reading declined sharply. The same trend was observed when the force reading was set to zero with UV-light switched on: switching it off led to a decrease in measured force, switching it back on to an increase in force (On-Off-On). These results indicate that the µFDS set-up is sufficiently sensitive to measure changes in muscle tone and stiffening following ChR2 mediated muscle contraction (see Suppl Fig 4).

**Figure 5.**
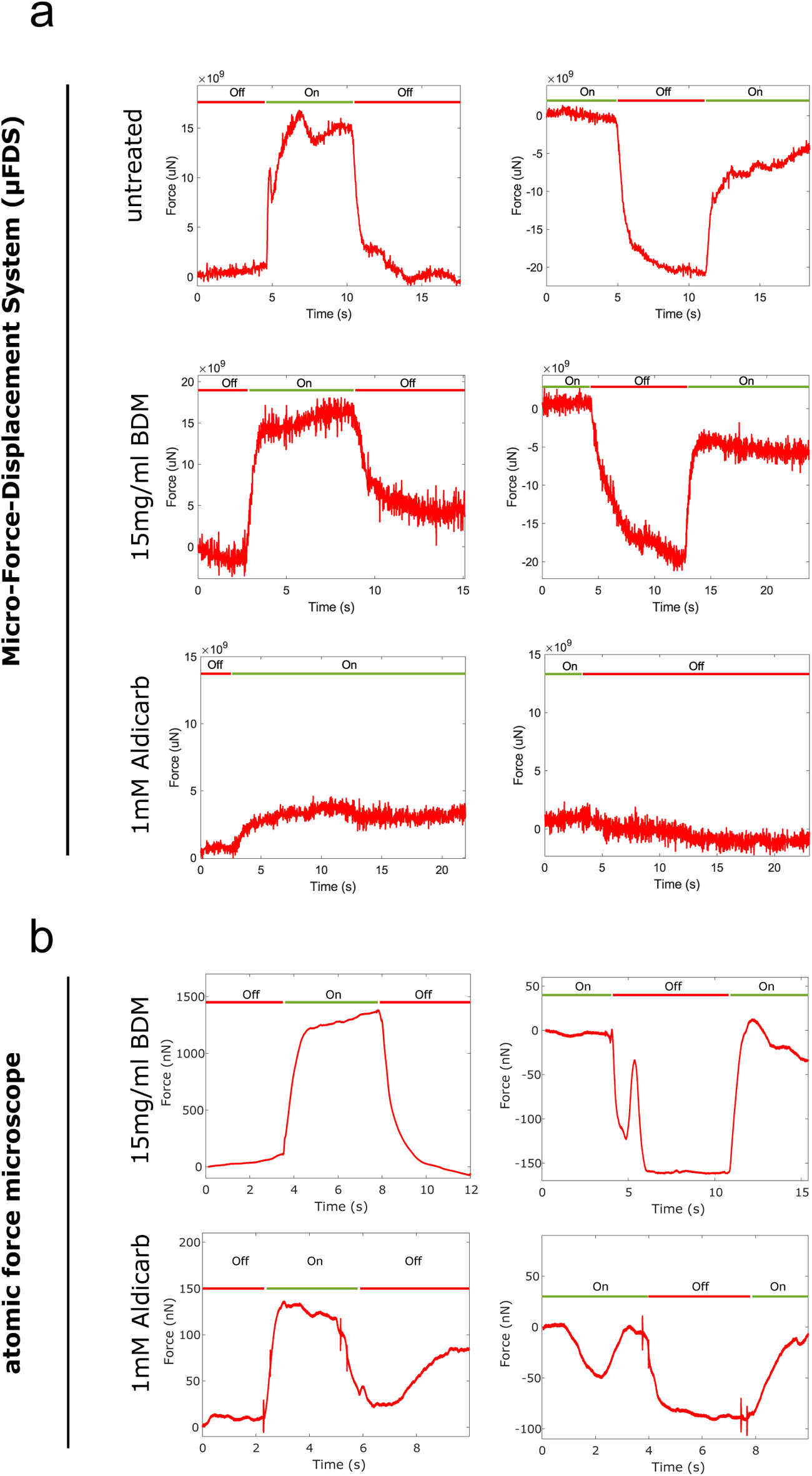
Force measured at a fixed indentation for indentation-clamped transgenic worms expressing ChR2 in muscle tissue. (**a**) µFDS measurements for untreated, BDM or aldicarb treated worms (light off: Off (*red*); light on: On (*green*)). (**b**) AFM measurements for BDM or aldicarb treated worms.

Next, we investigated whether worms treated with aldicarb, or BDM retained the ability to contract their muscles when under ChR2-induced muscle activation. Assuming BDM immobilises worms by relaxing the muscles, channel rhodopsin activation should still lead to muscle contraction and hence stiffening of the worm. On the contrary, aldicarb-treated worms (with hypercontracted muscles) would likely not increase their stiffness following ChR2 activation. Using both the µFDS (Fig 5a) or AFM (Fig 5b) set-up, BDM-treated worms showed a robust increase in stiffness upon ChR2 activation in both activation–deactivation sequences (On–Off–On or Off–On–Off). Untreated transgenic worms had responded similarly to light activation (Fig 5a). Hence it is possible to induce and to measure muscle contraction in worms treated with BDM, suggesting BDM leaves muscles in a relaxed or responsive state. Using the µFDS set-up no significant stiffness increase was detected for aldicarb-treated worms following UV-light exposure (Fig 5a and Suppl Fig 4a). Using the AFM set-up, however, we observed an increase in measured compressive force for approximately half of the worms following UV-light exposure (Fig 5b and Suppl Fig 4b). This suggests that responsive aldicarb-treated worms had contractile muscles which were not hypercontracted. For those worms that did not respond, muscles were either hyper-contracted already and unable to contract further or were unresponsive due to additional effects of aldicarb. We suggest the latter as the most likely explanation, as ChR2 activation-induced muscle contraction results in a stiffening of the worm whereas aldicarb-treated worms have reduced stiffness compared to BDM-treated worms (Fig 3).

## 4 DISCUSSION

### 4.1 FE model

In this study we generated new empirical data of small-scale indentations in *C. elegans* of up to 2 *µ*m, which we used subsequently to build a new *in silico* model to simulate the biomechanics of the worm. In distinction to previously published work, our model can permit to assigning Young’s moduli to three distinct compartments of the *C. elegans* (cuticle, muscle and pseudocoelom) based on the shape of experimentally acquired F-D curves. AFM data indicated that the F-D relation is highly non-linear (see for example Fig 1c), hence, pertinent methodological considerations in the *in silico* model are required. Our model differs from that of previous works, such as the “homogenous cylindrical shell with internal hydrostatic pressure” model by Park et al. (2007) and the “pressurized cylinder composed of two layers” model by Elmi et al. (2017), both of which considered *C. elegans* tissue mechanical behaviour as linear elastic. Our model effectively encompasses the nonlinearities stemming from the contact between the indenting device and the worm’s cuticle, and the inherent nonlinear biomechanical behaviour of *C. elegans*’ tissue.

Using experimental data to inform and tune the parameters of our *in silico* model, we successfully reproduced the experimentally F-D curves by varying the Young’s moduli of the different compartments. We were then able to evaluate the contribution of different compartments, and the effect of neuromuscular agents and chemicals on tissue biomechanics and *C. elegans*’ bulk stiffness. The present *in silico* model of the *C. elegans* comes however with simplifications with respect to (*a*) the idealized geometrical representation of the worm’s body, (*b*) the dermis and muscle tissue anisotropic biomechanical behaviour, (*c*) the transport of biofluids in the pseudocoelom, and (*d*) the swelling of its tissue due to biochemical factors. To increase the resolution and complexity of the model, new data from experimental modalities would be required which were not available during this work. This could include tensile testing of *ex vivo* specimen cut-outs of the cuticle at different orientations to measure anisotropy and employ optical tomography to measure fluid flow in the gonad and intestinal cavity of the worm.

### 4.2 Hydrostatic pressure

Our *in silico* modelling data suggest the pseudocoelom has a substantial impact on *C. elegans*’ overall biomechanics. Reduction of internal pressure by high salt exposure reduced the Young’s modulus of the pseudocoelom by 91% according to our model. This observation contrasts with the findings of Park et al. (2007) in two ways. Firstly, their study proposed that the mechanics of the ‘outer shell’ was the major contributor to worm stiffness, and secondly, that hyperosmotic stress increased worm stiffness. How can a deeper compartment like the pseudocoelom affect stiffness? The contribution of an underlying tissue compartment even if not directly indented, in our case the indentation depth reached by the AFM was maximum 1 *µ*m, is relevant to due to the model’s assumed incompressibility of all three compartments. As a result, the deformation of the outer compartment (cuticle) is transferred rigidly to the compartments below. To demonstrate the influence of the pseudocoelom on the F-D curve we decreased the Young’s modulus of the muscle compartment instead of the pseudocoelom. The simulated F-D curve no longer matched the experimental results, being shallower for smaller displacements due to the soft muscle compartment, and steeper for larger displacements due to the stiff pseudocoelom compartment. Secondly, Park and colleagues reported that hyperosmotic shock increased body stiffness due to salt-induced muscle contraction, meaning the reduced internal pressure had lesser impact on the overall body stiffness. In our experiments, exposure to a high salt concentration led to reduced stiffness. A key difference between the two studies is that in addition to salt, worms were treated with BDM for complete immobilization during our AFM measurements. BDM, acting as a muscle relaxant may have interfered potentially with any muscle contraction promoted by hyperosmotic conditions. It is worth noting however that the body wall muscles in BDM-treated worms remained responsive to induced contraction, as shown by ChR2 activation in the presence of BDM (Fig 5). However, it is possible that the pathway leading to hyperosmotic-induced muscle contraction may act upstream of BDM, and consequently, our measurement results primarily reflected the effect of a reduction in hydrostatic pressure. Nevertheless, our findings are consistent with the observation that a decrease in internal pressure due to cuticle puncture similarly reduces body stiffness (Park et al., 2007).

### 4.3 Aldicarb on stiffness and morphology

The neuromuscular agent aldicarb has been an invaluable tool in *C. elegans* research to investigate neuromuscular function and behaviour. By assessing mobility and paralysis in response to this agent, researchers have gained insights into neuromuscular and synaptic functions (Miller et al., 1996; Sieburth et al., 2005). A large body of work has linked sensitivity to these agents, resulting in paralysis and shrinkage, to hypercontraction of body wall muscles. To understand the contribution of the body wall muscles to overall stiffness of a worm, we modulated the muscle tone using aldicarb, expecting a stiffening due to muscle hyper-contraction. However, using two different force-indentation systems, we observed that administration of aldicarb reduces worm stiffness. The question is therefore whether the effect of aldicarb in addition to muscle contraction could also affect other properties of *C. elegans* and, if so, what would these be?

In our optogenetics experiment, aldicarb-treated worms either increased in stiffness or showed no response upon UV-light induced muscle contraction. Aldicarb-treated worms shrink, and they are softer than control worms (BDM-treated). In addition, our biomechanical model suggests the most substantial change in stiffness arises from the pseudocoelom compartment. We therefore propose that aldicarb-treatment ultimately decreases the *C. elegans* internal pressure. This reduction might be a secondary effect following an initial muscle contraction, possibly to counteract the pressure generated during muscle contraction. Worms exposed to high salt and aldicarb decreased in stiffness and volume, although there was an observable difference in relation to width and length which might suggest different underlying causes.

We did not observe any evidence for a muscle hyper-contraction. Firstly, in our optogenetic experiment aldicarb-treated worms were at least partially stiffening under UV-light exposure suggesting the muscles were in a responsive, as opposed to hypercontracted, state. Secondly, when administering aldicarb to untreated worms in our µFDS system we observed a gradual softening of the worm (Suppl Fig 5). Thirdly, the aldicarb is expected to inhibit ACE, acetyl-choline-esterase, leading to the accumulation of the neurotransmitter acetylcholine (ACh), which acts on different ACh receptors including ion-gated excitatory nicotinic AChR, muscarinic AChR and possibly inhibitory anion-gated ion channels (Putrenko et al., 2005). A study by the Hobert lab (Pereira et al., 2015) describes that more than 50% of neurons in *C. elegans* (159 out of 302) are cholinergic including motor neurons, sensory neurons, and interneurons. Sensory neurons, including mechanosensory neurons, have been reported to be involved in osmoregulation, none of which is directly cholinergic (details available from the [WORMATLAS](https://www.wormatlas.org/neurons/Individual%20Neurons/Neuronframeset.html) database). Though one of them, the URX neuron, has direct contact to the pseudocoelomic fluid. Moreover, some of the sensory neurons’ direct downstream target interneurons are cholinergic and hence speculatively effected by aldicarb.

When comparing the effects of aldicarb administered on the plate and in solution, worms on plates underwent additional shrinkage, resulting in reduced length and volume (Suppl Fig 3a). This might be attributable to the behaviour of worms, which swim in solution and crawl on plates. Swimming worms showing diminished pharyngeal pumping in comparison to their counterparts that crawl on agar plates (Vidal-Gadea and Pierce-Shimomura, 2012). When administered on an agar plate aldicarb is likely to undergo both digestion and absorption, enabling it to reach more tissue, whereas worms in solution will experience reduced intestinal uptake. Although not significant, worms on plates tended to be softer than worms treated in aldicarb solution. Correlation between size and stiffness is not necessarily linear, a reduction in volume does not have to equate to a proportional decrease in stiffness, and that there must exist a limit to how soft a worm can become due to the reduced pressure in the pseudocoelom.

## 5 CONCLUSIONS

We have investigated the biomechanics of *C. elegans* using empirical data together with an *in silico* model, using the FE method, to characterize the mechanical properties of three different worm tissue compartments. Building our model in a multi-compartmental way allowed us to assign individual Young’s moduli for these three compartments, i.e., cuticle and hypodermis, muscle and pseudocoelom, based on the shape of the force-indentation curve. Using our *in silico* model, we dissected the impact of osmotic shock and the neuromuscular agents (i.e., aldicarb, tetramisol) on the mechanics of each tissue compartment. Despite the assigned hyper-contracting muscle action of aldicarb, treatment led to a softening of the worm accompanied by shrinkage, with the largest impact observed on the mechanics of the pseudocoelom compartment. Optogenetic-induced muscle contraction stiffened the worm, even under aldicarb treatment altogether, thus indicating additional effects of aldicarb on the *C. elegans*. While we demonstrated the capacity of our *in silico* model to probe the impact of drugs in muscle function and hydrostatic pressure, the model can be in the future extended to study the biomechanical impact of mutations that for example affect the cuticle or muscle function, or of pharmacologicals directed to improve muscle function, or collagen production. Furthermore, the model can be used to study the impact on specific tissues mechanics through processes like ageing, or the impact of diets and nutrients, e.g., certain lipids or sugar types.

## AUTHOR CONTRIBUTIONS

C.L.E. and M.E. designed and performed the experiments and analysed the experimental data. M.S. and V.P. supplied critical support for data acquisition and analysis. C.R., N.C. and V.V. built the *in silico* model. C.R. and N.C. carried out the simulations. C.L.E., M.E. and V.V. provided feedback about the simulations. C.L.E., C.R., M.E. and V.V. wrote the manuscript. M.A.S. and M.S. reviewed the manuscript. M.A.S. and M.S. secured funding.

## REFERENCES

Altun, Z. and Hall, D. (2023). Handbook of C. elegans Anatomy. WormAtlas

AzimiHashemi, N., Erbguth, K., Vogt, A., Riemensperger, T., Rauch, E., Woodmansee, D., et al. (2014). Synthetic retinal analogues modify the spectral and kinetic characteristics of microbial rhodopsin optogenetic tools. Nat. Commun. 5. doi:10.1038/ncomms6810

Backholm, M., Ryu, W. S., and Dalnoki-Veress, K. (2013). Viscoelastic properties of the nematode Caenorhabditis elegans, a self-similar, shear-thinning worm. Proceedings of the National Academy of Sciences 110, 4528–4533. doi:10.1073/pnas.1219965110

Brenner, S. (1974). The genetics of Caenorhabditis elegans. Genetics 77, 71–94. doi:10.1093/genetics/77.1.71

Butt, H.-J. and Jaschke, M. (1995). Calculation of thermal noise in atomic force microscopy. Nanotechnology 6, 1. doi:10.1088/0957-4484/6/1/001

Dandekar, K., Raju, B., and Srinivasan, M. (2003). 3-D Finite-Element Models of Human and Monkey Fingertips to Investigate the Mechanics of Tactile Sense. Journal of Biomechanical Engineering 125, 682–691. doi:10.1115/1.1613673

Du, Y., Stewart, P., Hill, N. A., Yin, H., Penta, R., Köry, J., et al. (2023). Nonlinear indentation of second-order hyperelastic materials. Journal of the Mechanics and Physics of Solids 171, 105139. doi:10.1016/j.jmps.2022.105139

Elmi, M., Pawar, V. M., Shaw, M., Wong, D., Zhan, H., and Srinivasan, M. A. (2017). Determining the biomechanics of touch sensation in C. elegans. Sci. Rep. 7. doi:10.1038/s41598-017-12190-0

Essmann, C., Martinez-Martinez, D., Pryor, R., Leung, K.-Y., Krishnan, K. B., Lui, P. P., et al. (2020). Mechanical properties measured by atomic force microscopy define health biomarkers in ageing C. elegans. Nat. Commun. 11. doi:10.1038/s41467-020-14785-0

Essmann, C. L., Elmi, M., Shaw, M., Anand, G. M., Pawar, V. M., and Srinivasan, M. A. (2017). In-vivo high resolution AFM topographic imaging of Caenorhabditis elegans reveals previously unreported surface structures of cuticle mutants. Nanomedicine: Nanotechnology, Biology and Medicine 13, 183–189. doi:10.1016/j.nano.2016.09.006

Gilpin, W., Uppaluri, S., and Brangwynne, C. P. (2015). Worms under Pressure: Bulk Mechanical Properties of C. elegans Are Independent of the Cuticle. Biophysical Journal 108, 1887–1898. doi:10.1016/j.bpj.2015.03.020

Glenn, C. F., Chow, D. K., David, L., Cooke, C. A., Gami, M. S., Iser, W. B., et al. (2004). Behavioral Deficits During Early Stages of Aging in Caenorhabditis elegans Result From Locomotory Deficits Possibly Linked to Muscle Frailty. The Journals of Gerontology: Series A 59, 1251–1260. doi:10.1093/gerona/59.12.1251

Higuchi, H. and Takemori, S. (1989). Butanedione Monoxime Suppresses Contraction and ATPase Activity of Rabbit Skeletal Muscle. The Journal of Biochemistry 105, 638–643. doi:10.1093/oxfordjournals.jbchem.a122717

Izquierdo, P. G., O’Connor, V., Green, A. C., Holden-Dye, L., and Tattersall, J. (2021). C. elegans pharyngeal pumping provides a whole organism bio-assay to investigate anti-cholinesterase intoxication and antidotes. NeuroToxicology 82, 50–62. doi:10.1016/j.neuro.2020.11.001

Javer, A., Ripoll-Sánchez, L., and Brown, A. E. (2018). Powerful and interpretable behavioural features for quantitative phenotyping of Caenorhabditis elegans. Philosophical Transactions of the Royal Society B: Biological Sciences 373, 20170375. doi:10.1098/rstb.2017.0375

Jung, J., Nakajima, M., Kojima, M., Ooe, K., and Fukuda, T. (2012). Microchip device for measurement of body volume of C. elegans as bioindicator application. J. Micro-Nano Mech. 7, 3–11. doi:10.1007/s12213-011-0036-7

Lackner, M. R., Nurrish, S. J., and Kaplan, J. M. (1999). Facilitation of Synaptic Transmission by EGL-30 Gqα and EGL-8 PLCβ: DAG Binding to UNC-13 Is Required to Stimulate Acetylcholine Release. Neuron 24, 335–346. doi:10.1016/S0896-6273(00)80848-X

Lewis, J., Wu, C.-H., Levine, J., and Berg, H. (1980). Levamisole-resitant mutants of the nematode Caenorhabditis elegans appear to lack pharmacological acetylcholine receptors. Neuroscience 5, 967–989. doi:10.1016/0306-4522(80)90180-3

Lints, R. and Hall, D. (2005). Handbook of C. elegans Male Anatomy. WormAtlas

Mahoney, T., Luo, S., and Nonet, M. (2006). Analysis of synaptic transmission in Caenorhabditis elegans using an aldicarb-sensitivity assay. Nat Protoc 1, 1772–1777. doi:10.1038/nprot.2006.281

Miller, K., Alfonso, A., Nguyen, M., Crowell, J., Johnson, C., and J.B., R. (1996). A genetic selection for Caenorhabditis elegans synaptic transmission mutants. Proceedings of the National Academy of Sciences 93, 12593–12598. doi:10.1073/pnas.93.22.12593

Moore, B., Jordan, J., and Baugh, L. (2013). WormSizer: High-throughput Analysis of Nematode Size and Shape. PLOS ONE 8, 1–13. doi:10.1371/journal.pone.0057142

Mulcahy, B., Holden-Dye, L., and O’Connor, V. (2013). Pharmacological assays reveal age-related changes in synaptic transmission at the Caenorhabditis elegans neuromuscular junction that are modified by reduced insulin signalling. Journal of Experimental Biology 216, 492–501. doi:10.1242/jeb.068734

Nagel, G., Brauner, M., Liewald, J., Adeishvili, N., Bamberg, E., and Gottschalk, A. (2005). Light Activation of Channelrhodopsin-2 in Excitable Cells of Caenorhabditis elegans Triggers Rapid Behavioral Responses. Current Biology 15, 2279–2284. doi:10.1016/j.cub.2005.11.032

Nakajima, M., Ahmad, M., Kojima, S., Homma, M., and Fukuda, T. (2009). Local stiffness measurements of C. elegans by buckling nanoprobes inside an Environmental SEM. In 2009 IEEE/RSJ International Conference on Intelligent Robots and Systems. 4695–4700. doi:10.1109/IROS.2009.5354091

Nurrish, S., Ségalat, L., and Kaplan, J. M. (1999). Serotonin Inhibition of Synaptic Transmission: Gαo Decreases the Abundance of UNC-13 at Release Sites. Neuron 24, 231–242. doi:10.1016/S0896-6273(00)80835-1.

Opperman, C. and Chang, S. (1991). Effects of Aldicarb and Fenamiphos on Acetycholinesterase and Motility of Caenorhabditis elegans. J Nematol 23, 20–27. doi:PMID:19283090

Park, S.-J., Goodman, M., and Pruitt, B. (2005). Measurement of mechanical properties of caenorhabditis elegans with a piezoresistive microcantilever system. In 2005 3rd IEEE/EMBS Special Topic Conference on Microtechnology in Medicine and Biology. 400–403. doi:10.1109/MMB.2005.1548488

Park, S.-J., Goodman, M., and Pruitt, B. (2007). Analysis of nematode mechanics by piezoresistive displacement clamp. Proceedings of the National Academy of Sciences 104, 17376–17381. doi:10.1073/pnas.0702138104

Pereira, L., Kratsios, P., Serrano-Saiz, E., Sheftel, H., Mayo, A., Hall, D., et al. (2015). A cellular and regulatory map of the cholinergic nervous system of *C. elegans*. eLife 4, e12432. doi:10.7554/eLife.12432.

Petzold, B. C., Park, S.-J., Ponce, P., Roozeboom, C., Powell, C., Goodman, M., et al. (2011). Caenorhabditis elegans Body Mechanics Are Regulated by Body Wall Muscle Tone. Biophysical Journal 100, 1977–1985. doi:10.1016/j.bpj.2011.02.035

Putrenko, I., Zakikhani, M., and Dent, J. (2005). A Family of Acetylcholine-gated Chloride Channel Subunits in Caenorhabditis elegans. Journal of Biological Chemistry 280, 6392–6398. doi:10.1074/jbc.M412644200

Rahimi, M., Sohrabi, S., and Murphy, C. T. (2022). Novel elasticity measurements reveal C. elegans cuticle stiffens with age and in a long-lived mutant. Biophysical Journal 121, 515–524. doi:10.1016/j.bpj.2022.01.013

Sanzeni, A., Katta, S., Petzold, B., Pruitt, B., Goodman, M., and Vergassola, M. (2019). Somatosensory neurons integrate the geometry of skin deformation and mechanotransduction channels to shape touch sensing. eLife 8, e43226. doi:10.7554/eLife.43226

Schultheis, C., Liewald, J., Bamberg, E., Nagel, G., and Gottschalk, A. (2011). Optogenetic Long-Term Manipulation of Behavior and Animal Development. PLOS ONE 6, 1–9. doi:10.1371/journal.pone.0018766.

Sieburth, D., Ch’ng, Q., Dybbs, M., Tavazoie, M., Kennedy, S., Wang, D., et al. (2005). Systematic analysis of genes required for synapse structure and function. Nature 436, 510–517. doi:10.1038/nature03809

[Dataset] SIMULIA (2014). Abaqus 6.14 Documentation

Sneddon, I. N. (1965). The relation between load and penetration in the axisymmetric boussinesq problem for a punch of arbitrary profile. International Journal of Engineering Science 3, 47–57. doi:10.1016/0020-7225(65)90019-4

Soeno, Y., Shimada, Y., and Obinata, T. (1999). BDM (2,3-butanedione monoxime), an inhibitor of myosin-actin interaction, suppresses myofibrillogenesis in skeletal muscle cells in culture. Cell Tissue Res. 295. doi:10.1007/s004410051237

Stroustrup, N., Ulmschneider, B. E., Nash, Z. M., Isaac F., López-Moyado, J. A., and Fontana, W. (2013). The Caenorhabditis elegans Lifespan Machine. Nature Methods 10, 665–670. doi:10.1038/nmeth.2475

Vidal-Gadea, A. G. and Pierce-Shimomura, J. T. (2012). Conserved role of dopamine in the modulation of behavior. Communicative & Integrative Biology 5, 440–447. doi:10.4161/worm.19148

Wills, I. (2019). The Edisonian Method: Trial and Error (Springer International Publishing). 203–222. doi:10.1007/978-3-030-29940-8 10

Wolkow, C. and Hall, D. (2016). Handbook of C. elegans Dauer Anatomy. WormAtlas

Yemini, E., Jucikas, T., Grundy, L. J., Brown, A. E., and Schafer, W. R. (2013). A database of Caenorhabditis elegans behavioral phenotypes. Nature Methods 10, 877–879. doi:10.1038/nmeth.2560

