## Supplementary figures and images for "The influence of internal pressure and neuromuscular agents on *C. elegans* biomechanics: an empirical and multi-compartmental *in silico* modelling study"

### Supplementary Fig 1

#### Supplementary Figure 1

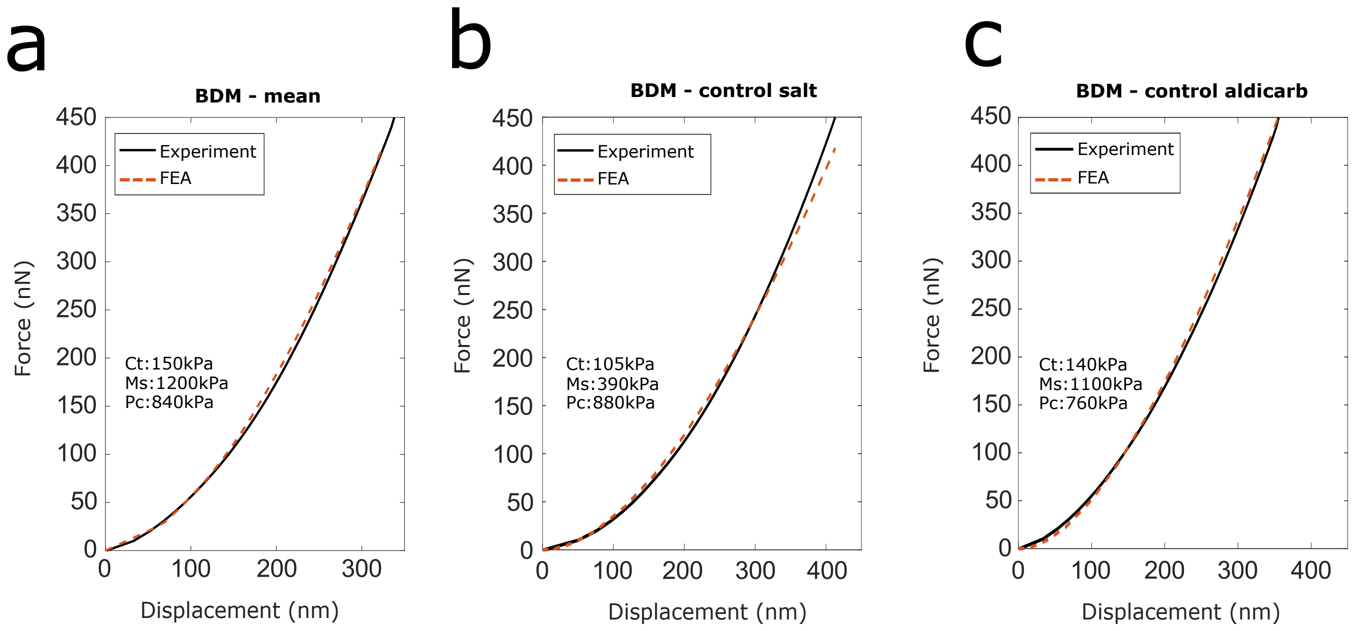

### Supplementary Fig 2

#### Supplementary Figure 2

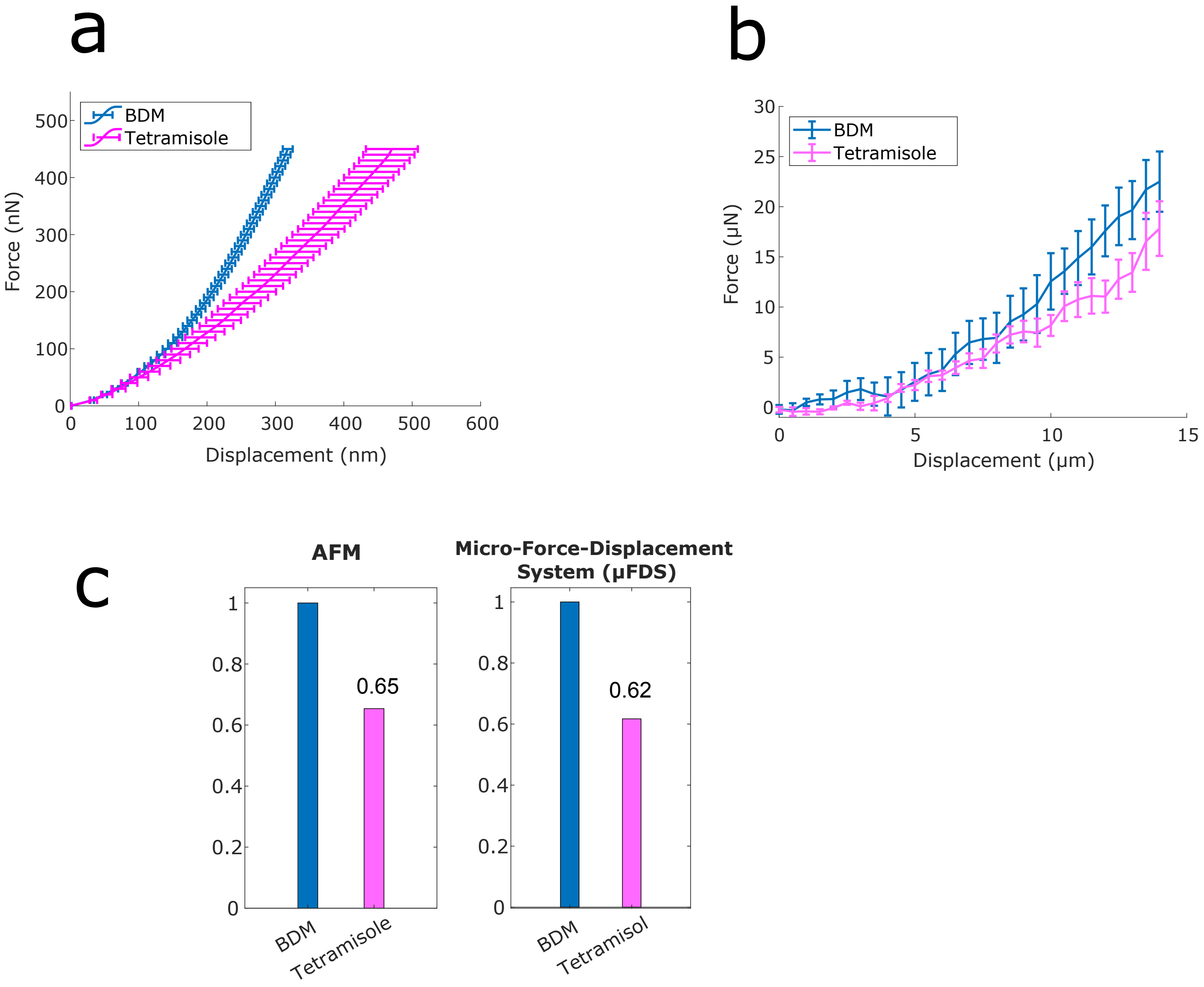

### Supplementary Fig 3

#### Supplementary Figure 3

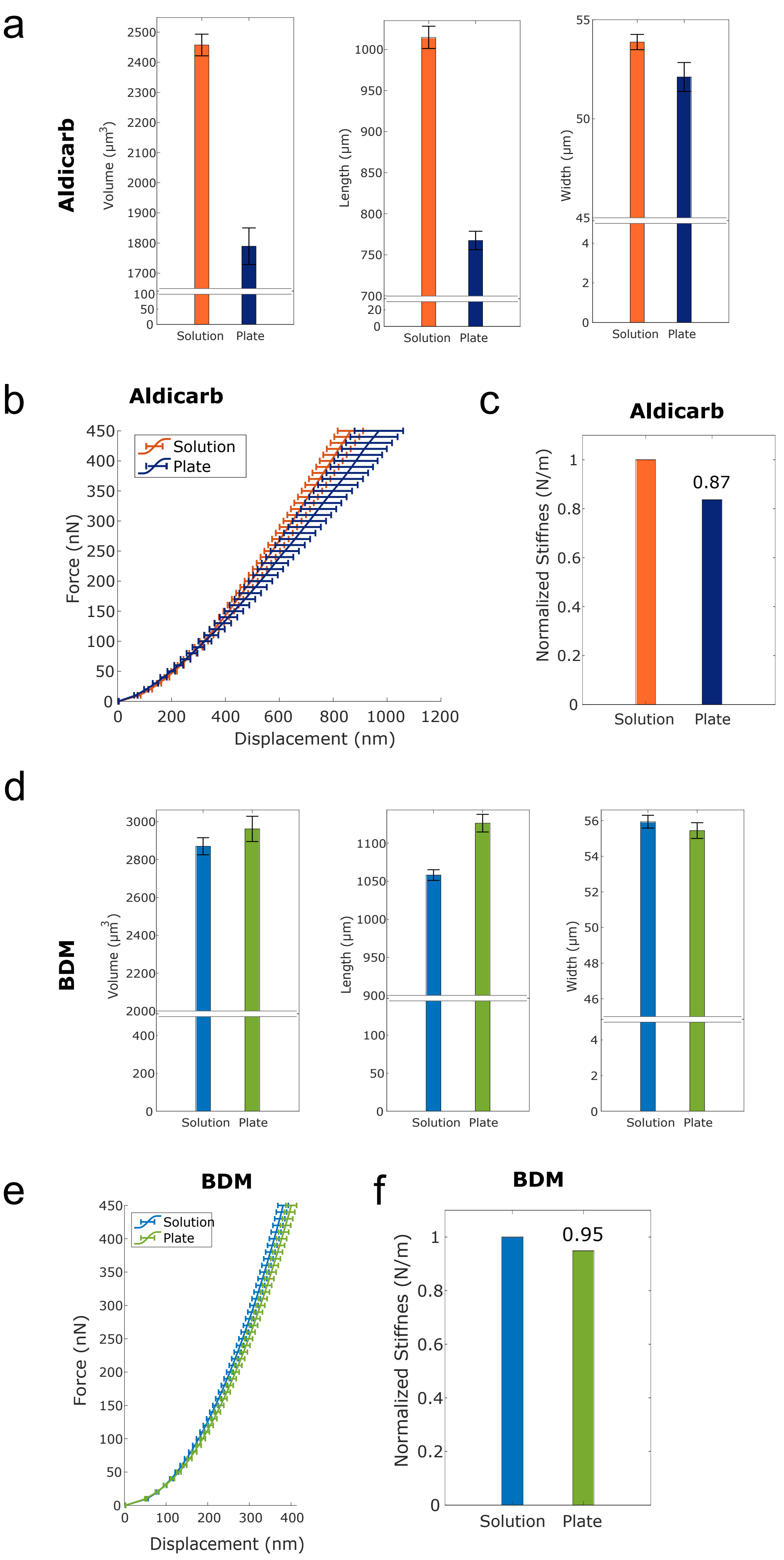

### Supplementary Fig 4

#### Supplementary Figure 4

a

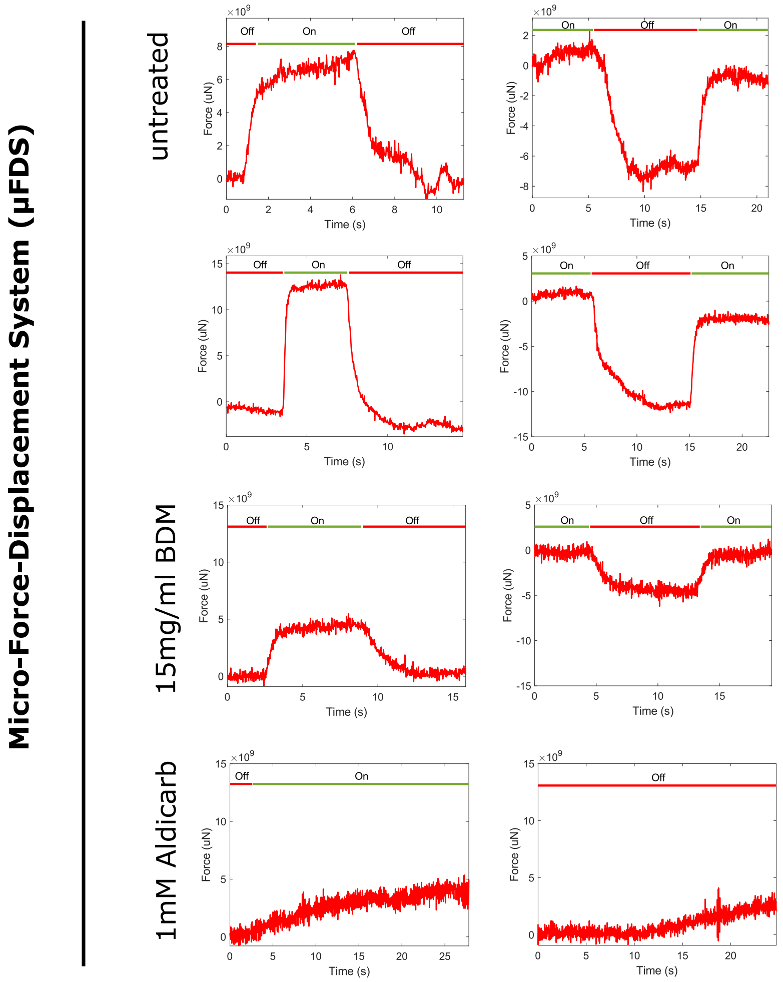


b

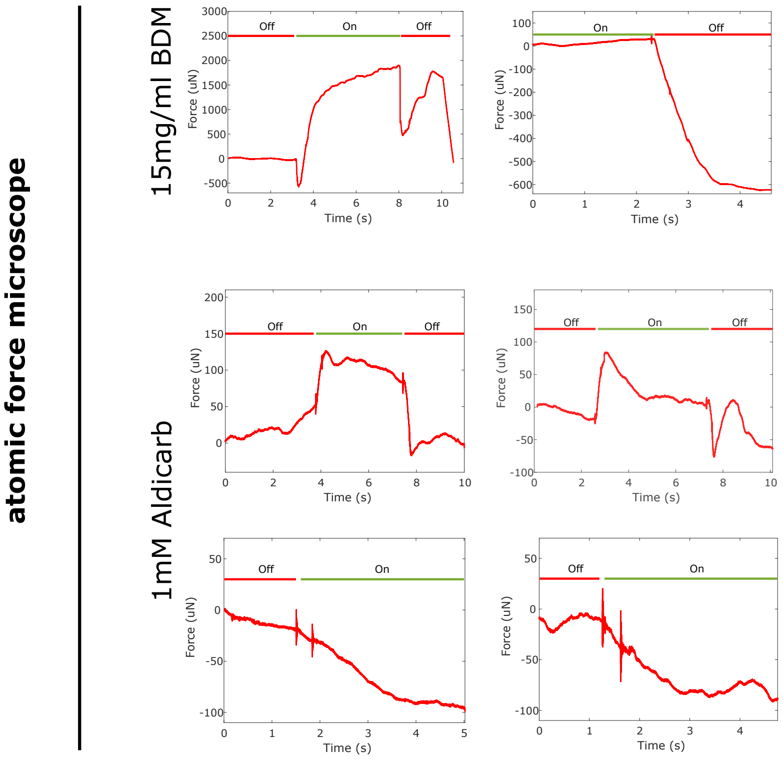

### Supplementary Fig 5

Supplementary Figure 5

a

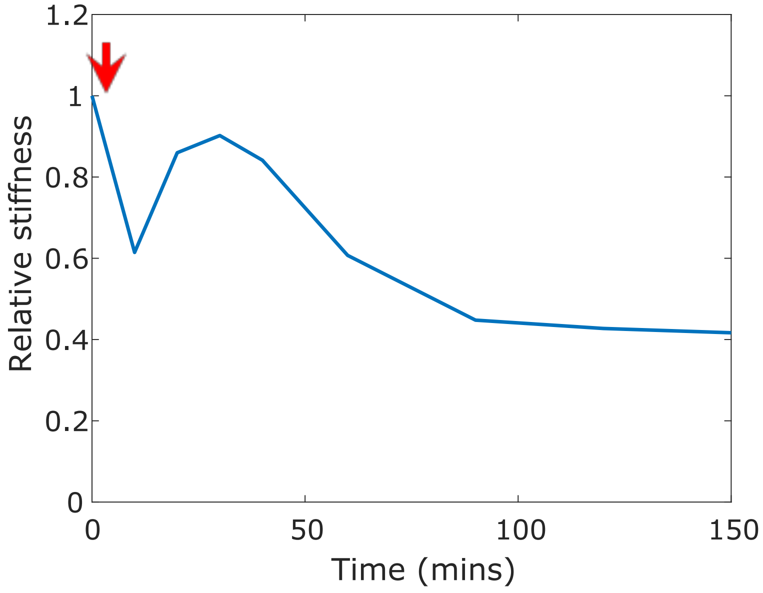


b

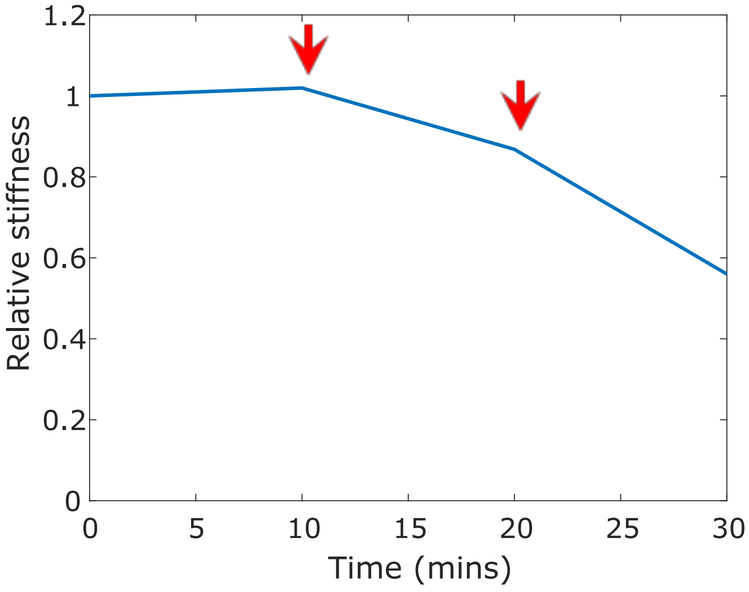
