## Supplementary Text 1 for "The influence of internal pressure and neuromuscular agents on *C. elegans* biomechanics: an empirical and multi-compartmental *in silico* modelling study"

### The influence of internal pressure and neuromuscular agents on *C. elegans* biomechanics: an empirical and multi-compartmental *in silico* modelling study

C.L. Essmann, M. Elmi, C. Rekatsinas, N. Chrysochoidis, M. Shaw, V. Pawar, M.A. Srinivasan, V. Vavourakis

### Finite element model of the *C. elegans*

A high-fidelity finite element (FE) model was developed to simulate the biomechanical behaviour of *Caenorhabditis elegans* (*C. elegans*) the key aspects of the model are documented here. The FE model was built using commercial software ABAQUS/Standard solver.[1] The FE model is three-dimensional in such that it is capable to simulate the whole body of the nematode – in this study we focus modelling the part of the body of interest that is tested from the AFM (see **Fig 1b** in the manuscript) or the μFDS (see **Fig 3b** in the manuscript) probe – with its tissue being described following a continuum-based approach, while mechanical interactions of the *C. elegans* with other objects (testing dish, agarose pad, indenter probe) have been explicitly introduced in the FE model using pertinent contact mechanics assumptions. In the next paragraphs, and following ABAQUS modules'[2] line-up, we provide technical description of the three-dimensional model of the worm (i.e., geometry, tissue regions), the FE mesh discretization and mesh convergence study considerations, the boundary conditions (BCs) and symmetry conditions for model simplification, and the justification of the nonlinear constitutive equation used to model *C. elegans* biomechanical behaviour.

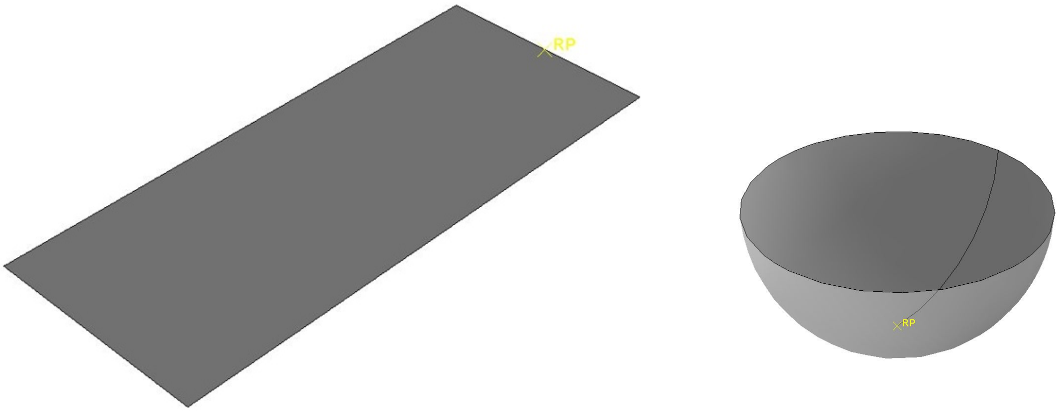


A B

**Figure 1.** (A) Horizontal glass surface and a reference point 'RP' assigned in ABAQUS' Part module, and (B) spherical surface of the indenter with second reference point 'RP' assigned at the tip of the probe.

#### *Geometry and material properties*

Following the experimental setup depicted in **Fig 1b** and **Fig 3b** in the manuscript, the *C. elegans* worm was positioned on a hydrated agarose gel and fixed laterally on the glass surface. Subsequently, the tip of the AFM probe moved normal towards the worm longitudinal axis to indent the skin. The present FE model was designed to simulate the experiment, evaluate the stress and deformations in *C. elegans*' tissue, and calculate the reaction forces (as they develop at the tip of indenter) while the indenter displaces towards and pushes the skin of the worm. It is very important

---

[1] ABAQUS. Abaqus 6.14 Documentation. Abaqus 614 Analysis User's Guide 2014.

[2] https://abaqus-docs.mit.edu/2017/English/SIMACAECAERefMap/simacae-c-topwhatismodule.htm#simacae-c-topwhatismodule

to highlight that in all simulations the rate of indentation was assumed slow, enough to ignore inertial effects in this study.

In ABAQUS Part module, the individual parts, i.e., elastic (deformable) and rigid (undeformable) bodies of the simulation, were sketched in the software 3D environment. The lateral and horizontal parts (glass plate) in contact with the *C. elegans* were modelled as rigid bodies explicitly, and an arbitrary reference point (see 'RP' label in **Fig 1** above) was assigned to initialize in ABAQUS the loading path of the indented towards the worm. In ABAQUS, it is essential to assign appropriate reference points, as they tie with the parts of the model and, hence, all applied forces, displacements and boundaries conditions are associated with these points of reference – boundary conditions are outlined in the next paragraph.

The *C. elegans* body was drawn in ABAQUS Part module and was modelled as an idealized cylinder having 56 µm diameter, and 1 mm long. Bellow, **Fig 2** depicts a close-up of the internal layout structure of the nematode's body, where the skin is shown in red (uniform thickness 0.6 µm), the muscle tissue region is shown in blue (uniform thickness 1.7 µm), and the internal tissue region of the worm is shown in green (radius 26.3 µm). The FE model was subsequently subdivided into fidelity regions through the thickness of the cross-section and across the axial direction of the worm model. The purpose inserting the fidelity zones was to allow in ABAQUS Mesh module to customize the FE mesh (refer to the "Finite element mesh generation" paragraph below) and optimize the density of the finite element grid in order to reduce the computational cost of the simulations for the *C. elegans* FE model.

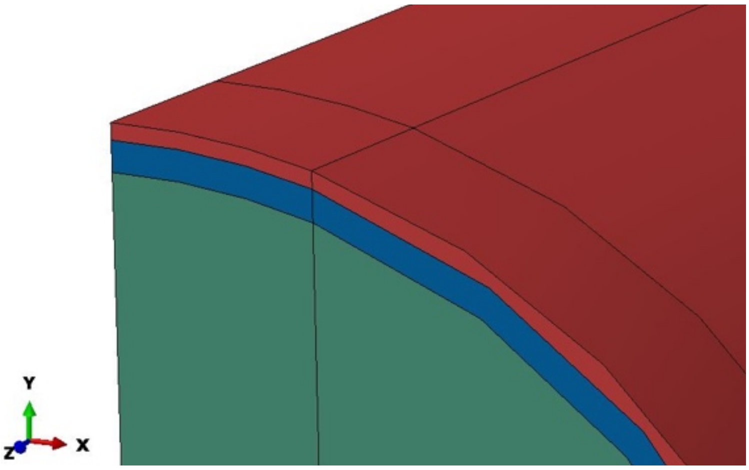


**Figure 2.** Cut through of the *C. elegans* geometry, as shown in ABAQUS Part module, depicts the three tissue regions considered in the 3D nematode model (red: skin; blue: muscle; green: guts).

Then as a second step, following ABAQUS Property module, we introduced the material definitions and assigned them to the appropriate regions of the geometry parts defined above. In this work, the mechanical behaviour of *C. elegans* tissue was modelled as a hyperelastic nonlinear material, which is typically used in biomechanical modelling of soft biological tissue (e.g., cardiovascular tissue, skin, muscle, and adipose tissue).

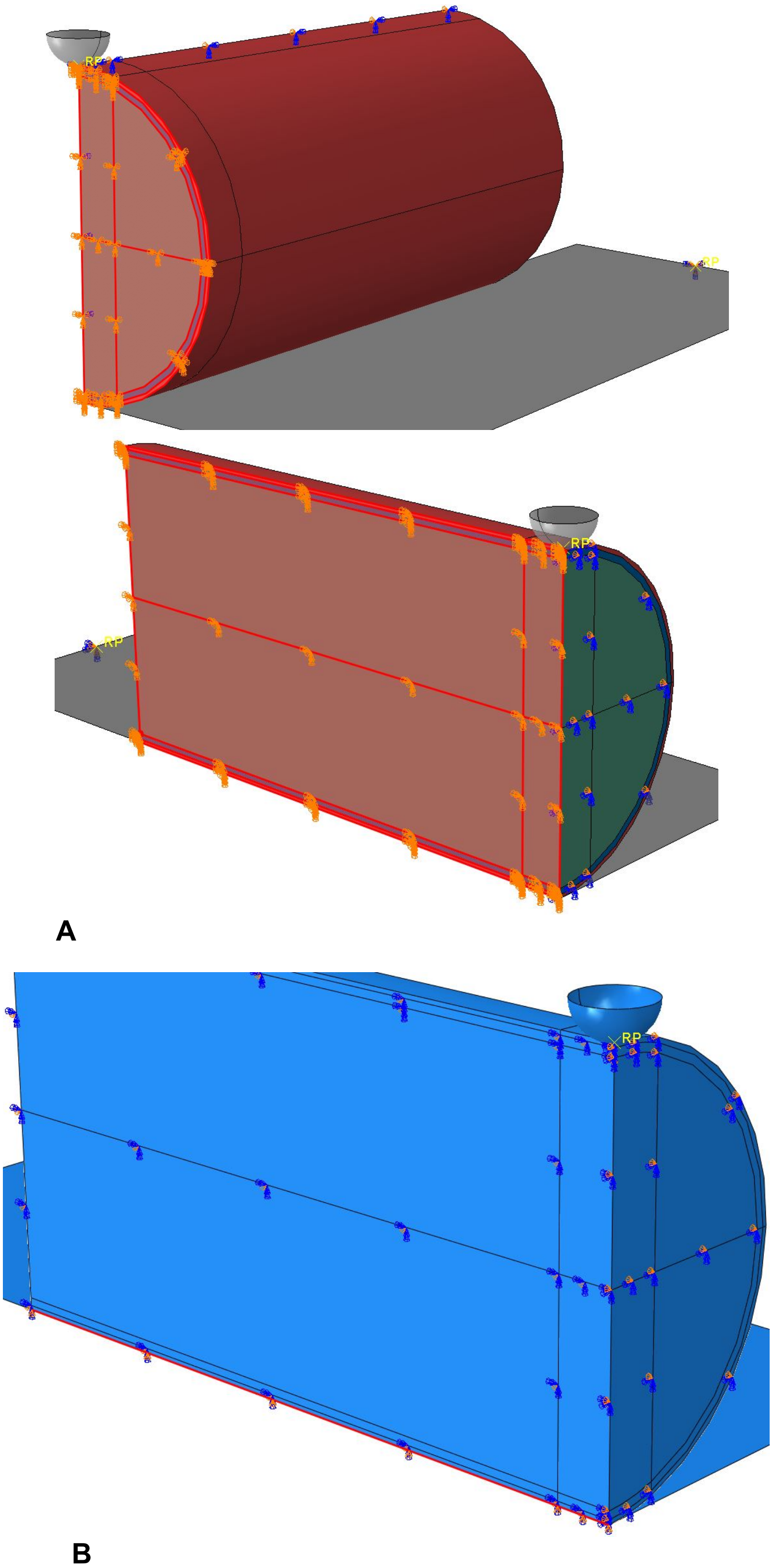


**Figure 3.** A: Views of the *C. elegans* geometry, as shown in ABAQUS Load module, illustrate the one quarter of the 3D nematode model (bottom: different colours represent separate set of BCs assigned to the model; opaque hemisphere: tip of the indenter positioned above the worm). B: Model was clamped (see red line across the cylinder—horizontal surface intersection) to constraint the model.

We adopted the neo-Hookean constitutive model, the strain-energy potential function is described by the formula:[3] $\Psi = 0.5\mu\,(\mathrm{tr}(\boldsymbol{F}\cdot\boldsymbol{F}^T) - 3) + 0.5\,K\,(J-1)^2$ where $\boldsymbol{F}$ the deformation gradient tensor, $\mathrm{tr}(\boldsymbol{F}\cdot\boldsymbol{F}^T)$ the trace of the left Green-Cauchy deformation tensor, $\boldsymbol{b}$, and $J = \det(\boldsymbol{F})$ the determinant of tensor $\boldsymbol{F}$; evaluation of the Cauchy stress tensor is computed through equation: $\boldsymbol{\sigma} = J^{-1}\,\boldsymbol{F}\cdot{}^{\partial\Psi}/_{\partial\boldsymbol{F}}$ which, after substitution and derivation on $\Psi$, gives: $\boldsymbol{\sigma} = \mu\,J^{-1}\boldsymbol{b} + K(J-1)\,\boldsymbol{I}$, with $\boldsymbol{I}$ being the identity matrix. Material parameters $\mu$ and $K$ under small deformations are equivalent to the shear and bulk modulus respectively, and they relate to the Young's modulus ($E$) and Poisson ratio ($\nu$) through: $\mu = E/2(1+\nu)$ and $K = E/3(1-2\nu)$. To effectively introduce tissue near-incompressibility, the Poisson ratio for all tissue regions was fixed to 0.45, while the elasticity modulus $E$ was modulated to interrogate *C. elegans* tissue stiffness under physiological (control) and treated conditions. Thus, having fixed $\nu$ and varying $E$, material parameters $\mu$ and $K$ were assigned in each tissue region (skin, muscle, internal); however, the maximum calculated bulk modulus was set uniform for all tissue regions.

#### *Boundary conditions and contact*

The third step in the FE model build was to assign proper boundary conditions to the 3D geometry of the *C. elegans* as well assign the appropriate external surfaces as prospective surface for contact conditions. In view of the geometry symmetry (cylindrical body of the worm) and the symmetries associated with the positioning and indentation of the worm, proper symmetry boundary conditions were assumed in the FE model. Symmetry conditions allowed to reduce the size of the model by one quarter and, therefore, decrease the computational cost considerably compared to the full-3D model of the nematode's body, without compromising numerical solution accuracy – justification to this is provided in the paragraph "FE model assessment" below. To this end, it should be highlighted that the total reaction force is equal with four times to the reaction force calculated in ABAQUS at the indenter reference point, due to the double symmetry conditions described above.

As shown in **Fig 3A**, symmetry boundary conditions were considered on the XY plane (red surface) and YZ plane (green surface) respectively, thus, producing one quarter of the full 3D model. In ABAQUS Load module, the vertical component (with respect to the corresponding symmetry plane) of the displacement vector was set zero for each symmetry plane respectively, while across the intersection line of the cylindrical geometry (worm) and the horizontal rigid surface (glass) all displacement components were set zero, as shown in Figure 3B. This constraint was imposed to avoid potential rigid-body movement FE simulation solutions.

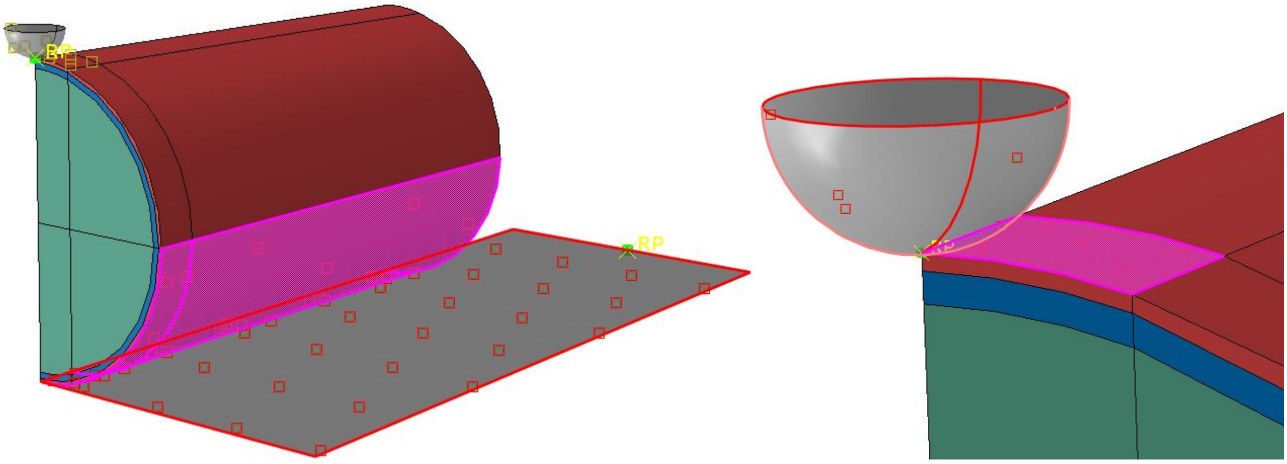


**Figure 4.** Surfaces marked for contact conditions in ABAQUS Interaction module (left: contact between the worm and the glass plate shown grey; right: detail of the contact between the worm and the tip of the indenter.

[3] K.-J. Bathe. Finite Element Procedures. ISBN-13: 978-0979004957

In addition to the conventional BCs, appropriate contact conditions were introduced in the FE model. As illustrated in **Fig 4**, we marked manually the surfaces of the *C. elegans* model that are anticipated during the simulation to come in contact (with self or other geometric parts of the ABAQUS model design). The surfaces marked pink in **Fig 4** above designate the 'slave' contact surfaces while the red-crossed line surfaces (the glass plate on the left and the tip of the indented on the right) designate the 'master' contact surfaces respectively. The convention, in ABAQUS and in most FE software, assumes a 'master' surface (or body) being the rigid (or hardest) part of the FE analysis – here the glass plate surface and the indenter tip was considered as such. The same convention assumes a 'slave' surface (or body) being the elastic (or softest compared to the 'master') part of the FE analysis. As outlined above, a reference point was assigned to it (see **Fig 1B** and **Fig 4**) to apply the sequential displacement increments to the indenter surface and model its vertical (rigid body) translation. ABAQUS Interaction module provides two different approaches to impose contact between geometrical entities: node-to-surface and surface-to-surface contact.[4] Both approaches were modelled and examined in our present analyses, the simulation results of which (not presented in this document) exhibited almost identical results – the surface-to-surface contact required less computational time. In addition, two modes of contact were tested: 'soft contact' (frictionless sliding between surfaces) and 'hard contact' (sliding between surfaces with friction).

Simulation trials – results not included in this supplementary material document – helped us to converge towards adopting the 'hard contact' option in this study. The 'hard contact' compared to the 'soft contact' was observed to recapitulate the deformations of the worm during indentation more realistically, while also the FE model exhibited computational stability. The present hard contact model assumed for Coulomb friction law with a coefficient of friction set to 0.01. To summarise, all simulation results shown in our paper consider ABAQUS' surface-to-surface contact approach and hard contact model.

#### *FE mesh generation*

Next step in the model preparation involves the FE mesh generation. The *C. elegans* model was processed in ABAQUS Mesh module to generate the finite elements that would fill the volume of the 3D model. In this work we enforced the generation of a hexa-dominant mesh (bilinear, 8-node hexahedral elements) with space varying density. As seen in **Fig 5**, the hexahedral elements were smaller at the contact surface, i.e., the fidelity region at the worm model right below the indenter, which was naturally extended to the fidelity regions adjacent to the contact surface one. Also, as evident in this illustration, for the skin and muscle tissue regions, at least two FEs were introduced across the layer thickness dimension to ensure numerical results quality (especially with respect to mechanical stresses predictions). More about the mesh convergence and FE stability is presented in the next paragraph, while more illustrations of the fidelity regions considered in the present model can be seen in **Fig 2** and **Fig 4** respectively.

[4] https://abaqus-docs.mit.edu/2017/English/SIMACAEITNRefMap/simaitn-c-contactoverview.htm

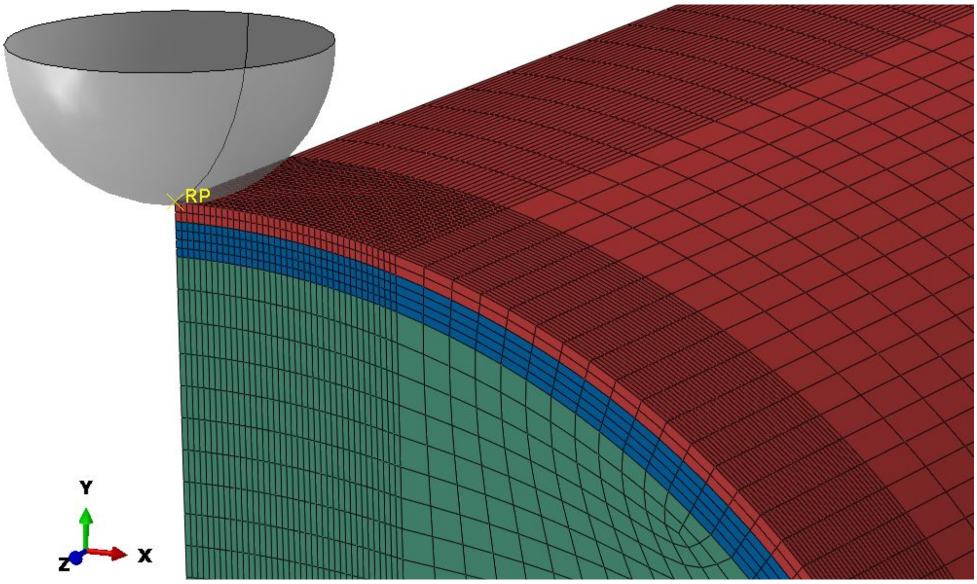


**Figure 5.** Close-up of the FE discretization of the *C. elegans* model (and tip of the indenter) that consists of hexahedral finite elements.

### *FE solver settings*

The problem under investigation considers nonlinear mechanics due to contact boundary conditions and the material model adopted to describe *C.elegans* biomechanics. In light of the low-rate of mechanical experimentation on the nematodes, the present study assumes a quasi-static loading and inertial loading effects are ignored. The optimum settings for the present problem which provide converged results (along with the respective mesh discretization), were set as follows. The (pseudo) time duration of the indentation simulation was set to 0.5 *time units*, while the (pseudo) time increments between sequential solution steps being automatically adapted by ABAQUS to ensure numerical convergence. The (pseudo) time increment was allowed to vary between $5 \cdot 10^{-7}$ and 0.05 *time units*, and the maximum number of (pseudo) time increments was set to 750 in ABAQUS' implicit solver.

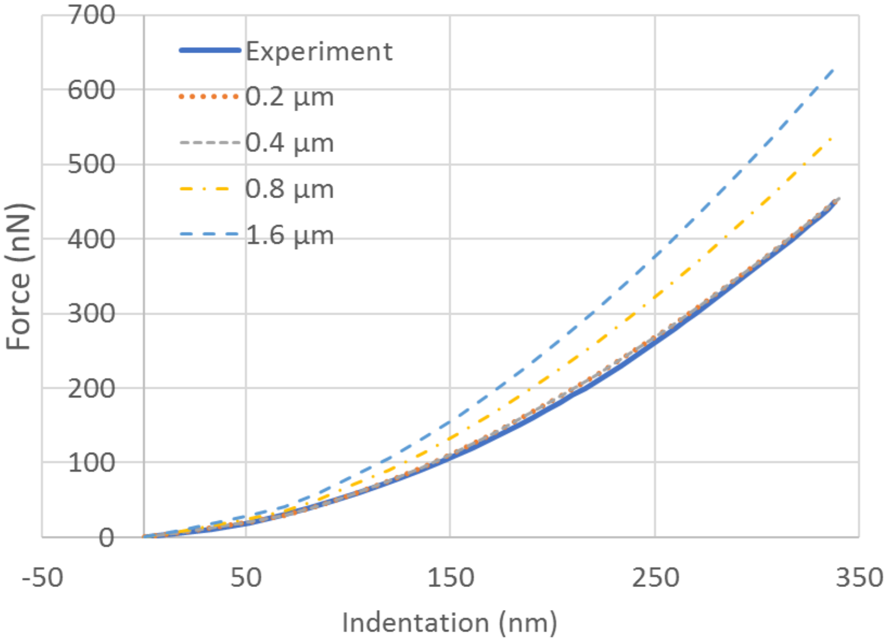


**Figure 6.** Force-displacement line plots compare the experimental data for the BDM/control case against the FE simulated data of the *C. elegans* FE model.

#### *FE model assessment*

Initially a FE convergence study was carried out to determine the optimal element length that would warrant accurate simulation results at an acceptable computational cost. As a reference for the convergence study, the data of the control scenario of the BDM case (see manuscript) was selected. Four different FE discretizations were considered the pattern of which was identical albeit the minimum edge length of a hexahedral element ranged from 0.2 μm up to 1.6 μm.

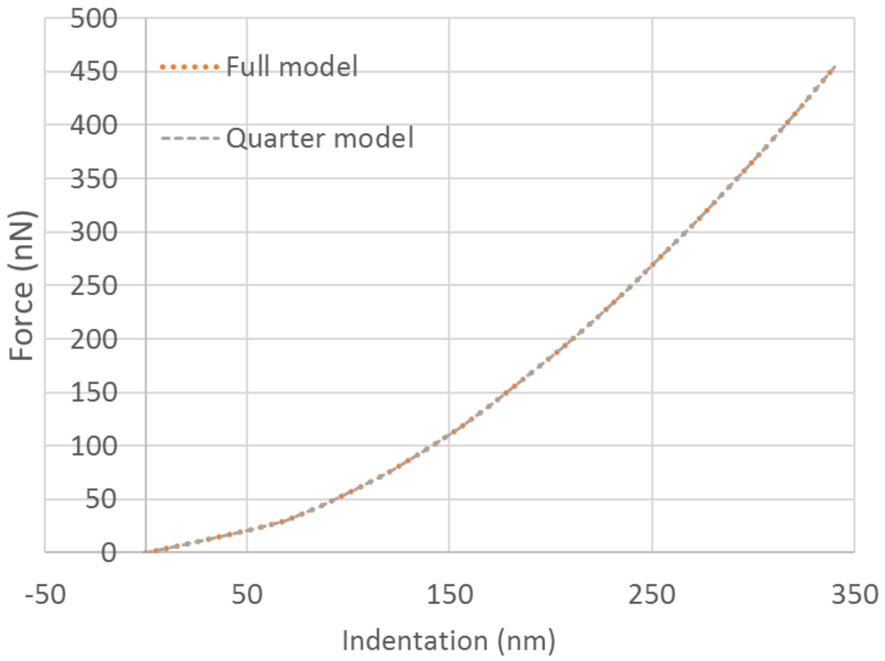


**Figure 7.** Force-displacement line plots compare the FE simulated data of the *C. elegans* model when considering the full 3D geometry model against the 'reduced size' model due to symmetry.

In **Fig 6**, the reaction force (experimentally measured and numerically calculated data) against the indentation depth at the tip of the indenter was plotted. As seen in this graph, the FE simulation results for 0.2 μm and 0.4 μm minimum edge size give very good agreement with the experimental results, while FE numerical solution demonstrates excellent convergence for when increasing the mesh density from a coarse mesh (minimum edge length 1.6 μm) to denser mesh densities (0.8 μm, 0.4 μm, and 0.2 μm) as expected.

Next, we compared the simulation results produced by the FE model when considering the full three-dimensional body of the *C. elegans* model against the simulation results of the FE model with symmetry BCs accounted in. As evidently shown in the line plot of **Fig 7**, the simulation results' agreement is excellent and confirms the validity of the symmetry conditions introduced in the *C. elegans* FE model.

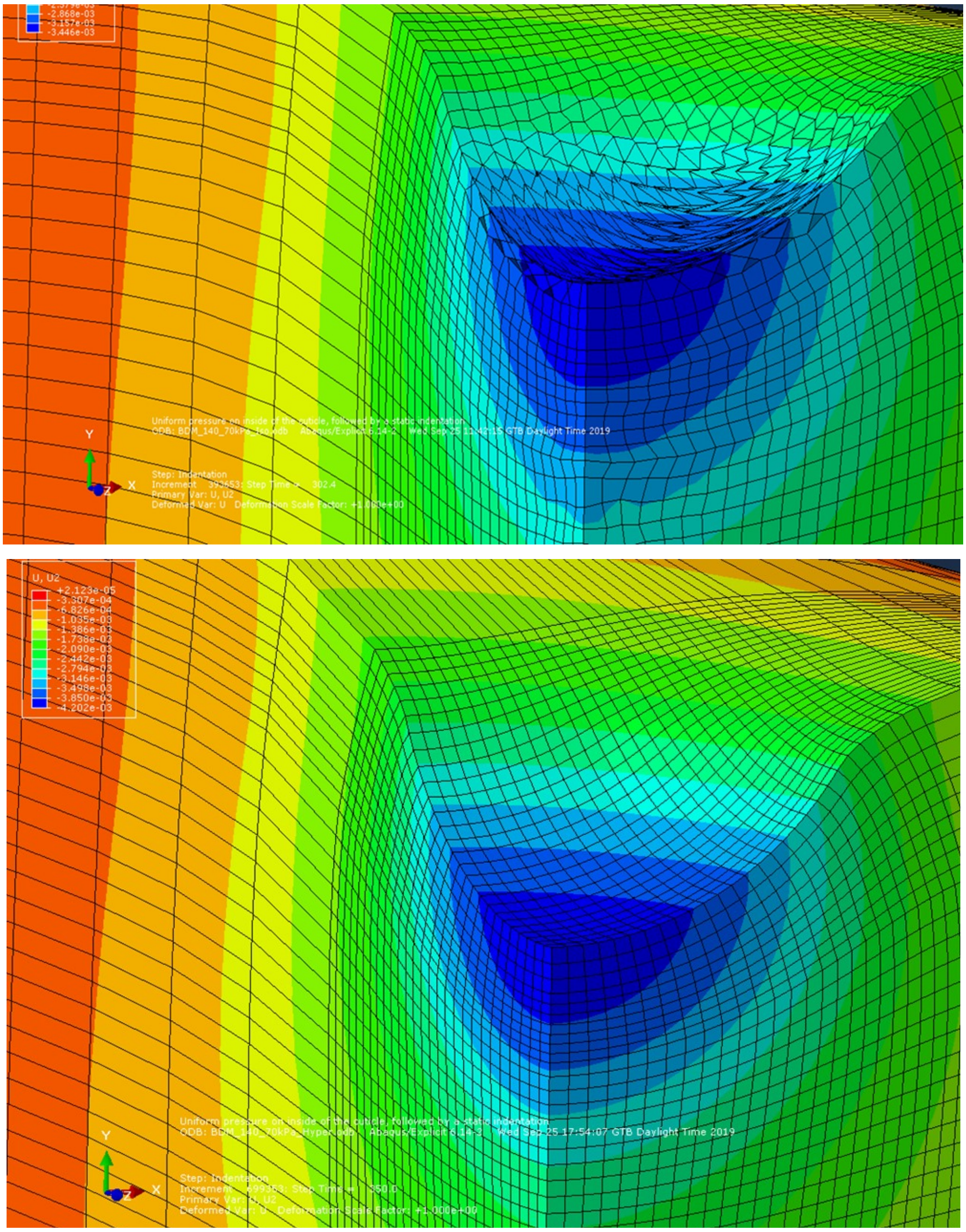


**Figure 8.** Vertical displacement contour plots of the FE simulations when (top) no numerical treatment was considered to overcome volumetric locking, whereas (bottom) numerical control of ‘hourglass effects’ in ABAQUS produces realistic simulation results.

Finally, we tested the FE model performance and capacity to manage numerical instabilities coming from volumetric locking,[5] also known as ‘hourglass effect’ – this is a typical numerical phenomenon attributed to material constraints or inaccurate numerical integration. In the present *C. elegans* FE model we anticipate volumetric locking to originate from the near- or incompressible behaviour of the nematode soft tissue biomechanics. In FEs, hourglass modes are nonphysical, zero-energy modes of deformation that produce zero strain (and thus no stress), and they occur only in under-integrated elements. To circumvent this, ABAQUS/Standard provides necessary controls to overcome ‘hourglass effects’ by which the stiffness coefficients are based on an enhanced strain method co-rotational formulation (no scale factor on the element stiffness is required). In **Fig 8** shown

[5] W.K. Liu, B. Moran, T. Belytschko, K. Elkhodary. Nonlinear Finite Elements for Continua and Structures. Wiley 2nd Edition, ISBN-13: 978-1118632703

above, the effect of hour-glassing in FEs locking is illustrated, while at the bottom of the same figure the simulation results demonstrate hourglass elimination.
