## Supplementary Table 1 for "The influence of internal pressure and neuromuscular agents on *C. elegans* biomechanics: an empirical and multi-compartmental *in silico* modelling study"

### The influence of internal pressure and neuromuscular agents on *C. elegans* biomechanics: an empirical and multi-compartmental *in silico* modelling study

C.L. Essmann, M. Elmi, C. Rekatsinas, N. Chrysochoidis, M. Shaw, V. Pawar,
M.A. Srinivasan, V. Vavourakis

Bulk stiffness maximum values of the *Caenorhabditis elegans* (*C. elegans*) that were used for the normalization of the data shown in the corresponding bar plot, see first column of the table.

| | |
|---|---|
| **Fig 2b** | 1.64 N/m |
| **Fig 3d** | 2 N/m (*left*) and 1.46 N/m (*right*) |
| **Supplementary Fig 1c** | 2.2 N/m (*left*) and 2 N/m (*right*) |
| **Supplementary Fig 2c** | 0.698 N/m |
| **Supplementary Fig 2f** | 2.24 N/m |
